# Hepatic fructose metabolism is antagonized by growth hormone/insulin-like growth factor signaling via regulation of ketohexokinase expression

**DOI:** 10.1101/2025.01.01.630846

**Authors:** Salaheldeen Elsaid, Prathibha Meesala, Youngshim Choi, Liqing Yu, Yang Xiao, Mitchell A. Lazar, Patrica Perez-Matute, Edward O. List, Lanuza AP Faccioli, Yiyue Sun, Rodrigo M Florentino, Alejandro Soto-Gutierrez, Jose G. Pichel, Darlene E. Berryman, Dirk Mayer, John J. Kopchick, Sui Seng Tee

## Abstract

Overconsumption of added sugars such as fructose has been associated with a remarkable decline in metabolic health. Fructose is primarily metabolized by the small intestines and the liver, via phosphorylation mediated by ketohexokinase (KHK). KHK activity is traditionally viewed as lacking negative feedback mechanisms, such as those present to limit glucose metabolism, leading to excessive fat accumulation characteristic of metabolic dysfunction-associated liver disease (MASLD). In this study, we observe KHK downregulation in hepatocytes of diet-induced and genetic models of MASLD. Reduced KHK coincides with decreased flux of fructose-derived carbons into glycolytic and amino acid metabolic pathways, suggesting the presence of mechanisms that limit KHK-mediated fructolysis in the liver. We subsequently focused on the growth hormone (GH)/insulin-like growth factor (IGF) signaling pathway as a potential mechanism antagonizing KHK expression. In transgenic mice with enhanced GH signaling, KHK levels are reduced, whereas reduced GH activity leads to increased KHK expression. Additionally, administration of GH and IGF-1 in liver cell cultures induces time-dependent degradation of KHK, facilitated by direct interactions between KHK and the IGF-1 receptor (IGF-1R). Single-nuclei RNA sequencing revealed elevated IGF-1R expression in hepatocytes from diet-induced MASLD mice, supported by human MASLD patient samples, which also show reduced KHK expression. Taken together, these findings describe a novel pathway by which GH/IGF-1 signaling regulates KHK, offering new insights into how the liver adapts to metabolic stress to limit fructose-driven liver dysfunction.

## Introduction

High fructose corn syrup (HFCS) consumption has increased by 33% in the last five decades ^1^. Excessive fructose intake in beverages and sweets has been linked to MASLD and its progressive stage nonalcoholic steatohepatitis (NASH) ^2,3^. This poses substantial public health challenges due to their rising incidence and the risk of progressing to cirrhosis and hepatocellular carcinoma (HCC)^4^. By 2050, it is estimated that approximately 24.3% of patients awaiting transplantation will have cirrhosis and HCC resulting from NASH^5^. Upon ingestion, 70% of ingested fructose is transported from the small intestine to the liver, where it undergoes metabolism^6^. The liver is the largest organ that metabolizes fructose due to the presence of the major fructolytic enzymes^6,7^. Unlike glucose metabolism, fructose metabolism lacks a negative feedback mechanism to regulate its breakdown, leading to continuous fructolysis ^2^. This uncontrolled fructolysis has been associated with the development of metabolic disorders such as cardiovascular disease^8^, insulin resistance, and type 2 diabetes ^9,10^. Hence, there is an unmet need to outline fructose metabolism and the molecular mechanism that links it to different morbidities, especially in the liver ^11^.

The heterogeneity and complexity of NASH has made it challenging to identify a single therapeutic target for its reversal ^12^. Over the past decade, numerous phase II/III clinical trials ^13^ have demonstrated encouraging outcomes. These drugs target different pathways, including lipid synthase^14,15^, Farnesoid X receptor signaling^16^, peroxisome proliferator-activated receptor signaling ^17^, hepatocellular injury, and inflammatory signaling^18^. Despite these advancements, the absence of established pharmaceutical agents specifically designed for NASH treatment persists. Thus, the identification of novel therapeutic targets remains an imperative and pressing objective within the research domain.

Fructose, identified as a key player in NASH^19^, also exhibits a notable correlation with fibrosis severity^20^. Chronic fructose intake has been linked with the development of MASLD and progressive stage NASH^2,21^. It imposes metabolic stress on the liver triggering de novo lipogenesis (DNL) and ATP depletion ^22,23^ as well as the production of harmful metabolites such as uric acid^24^. This results in multiple pathological consequences such as insulin and leptin resistance^25^, vascular inflammation^3^ and ultimately disrupting glucose homeostasis and contributing to the progression from prediabetes to type 2 diabetes^26^. Hence, it is important to understand the regulatory mechanisms governing fructose metabolism to mitigate its deleterious effect on the liver^11,27^.

Fructolysis involves several genes such as ChREBP, a key transcription factor that regulates genes associated with fructolysis, glycolysis, and lipogenesis^28–31^. It regulates transcription of genes including Glut5, fructokinase or ketohexokinase (KHK), Glut2, liver pyruvate kinase, acetyl CoA carboxylase, and fatty acid synthase (FASN). Targeting these genes could offer novel therapeutic interventions for NASH/NAFLD^32–34^.

KHK is an enzyme that is mainly expressed in the liver and that phosphorylates fructose into fructose-1-phosphate in a process lacking negative feedback^20^. Aldolase-B enzyme splits it into dihydroxyacetone phosphate and D-glyceraldehyde that contribute to lipogenesis^35^. Understanding the interaction between these key players in fructose metabolism can elucidate the mechanism by which fructose exerts its harmful effects on the liver.

In humans, KHK gene is located on chromosome 2p and comprises nine exons, with two exons (3a and 3c) mutually spliced to form two isoforms KHK-A and KHK-C^36,37^. The latter is the major phosphorylating enzyme due to its lower K_m_, indicating a higher affinity for fructose^36^. Notably, KHK-A exhibits a high K_m_ and lower affinity for phosphorylating fructose^36^. It exerts a protein kinase activity facilitating nucleotide synthesis through the polyol pathway, promoting epithelial to mesenchymal transition and metastasis, and activating the antioxidant pathway in cancer cells^38,39^. In hepatocellular carcinoma cells, transition in expression from KHK-C to KHK-A is mediated by hnRNPH1 and hnRNPH2 in a c-Myc-dependent mechanism^40^. The intriguing variation in KHK function and its distinct distribution highlight its crucial role in regulating various metabolic activities in both normal and cancer cells.

The growth hormone and insulin-like growth factor-1 (GH-IGF1) axis is emerging as a potential therapeutic target in NAFLD because of its lipolytic, anti-inflammatory, and anti-fibrotic properties^41–43^. Approximately 37 GH receptor knockouts have been studied in mice^44^. Notably, liver-specific growth hormone receptor knockout (GHR KO) have increased KHK expression ^45^. This increase can be partially reversed by infusion of insulin-like growth factor 1 (IGF1) and completely restored to normal levels by overexpression of STAT5^45^. Deletion of GHR in the liver has been associated with increased lipid accumulation and steatosis in mice ^46^ as well as in pigs^47^. Based on these findings, the present study aims to investigate the hormonal regulation of fructolytic activity by modulating KHK in liver cells. Understanding this mechanism may provide valuable insights into therapeutic strategies for managing MASLD by regulating fructose metabolism.

## Results

### 1- Hepatic KHK is downregulated in diet-induced models of NAFLD

Gubra-Amylin NASH or (GAN) diet is a high-fat (40% kcal from fat), high-fructose (22% kcal from fructose), and high-cholesterol (2% wt/wt) diet that induces obesity, insulin resistance, hepatic steatosis, inflammation, and fibrosis in mice, recapitulating the pathogenesis of human NASH^48,49^. Thirty wild type C57BL/6 male mice of 14 weeks age were divided into two groups. The first group was fed a standard diet (STD), while the second group received a GAN diet for 12 weeks (Fig.1). After this period, mice underwent extensive metabolomic studies. Finally, they were sacrificed, and their livers were collected for histological, immunohistochemical, and molecular analysis. Mice on the GAN diet showed significant body and liver weight gain and a modest, non-significant increase in blood glucose levels (Fig. 1-B).

**Fig (1).**
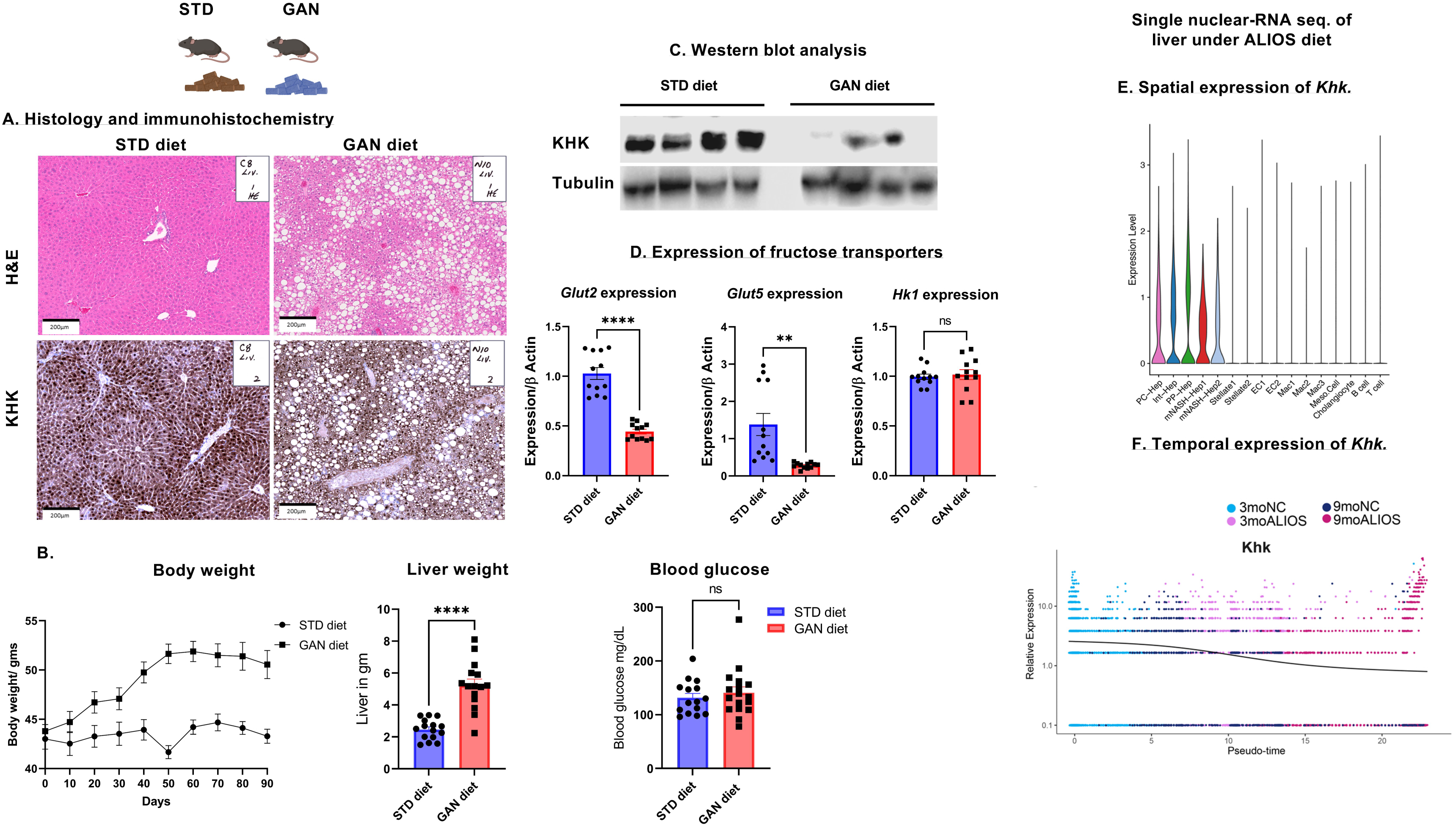
Khk downregulation was associated with induction of NASH. (A) Representative H&E micrographs showing macrovesicular and microvesicular steatosis, a hallmark of NASH. Immunohistochemical analysis reveals that in healthy liver tissue, KHK exhibits a periportal and interportal expression pattern. (B)Body weight was significantly increased in mice under GAN diet, the blood glucose level didn’t show significant difference. (C) Western blot analysis further confirms the decrease in KHK protein levels in NASH liver samples compared to healthy controls. (D)Quantification of KHK protein levels demonstrates a significant reduction in NASH liver tissue. (E) Gene expression analysis showed a significant reduction in fructose transporters in NASH liver. *Hk1* showed no marked change between the two groups. **(F)** Violin plots of differential *Khk* gene expression in mouse livers: *Khk* expression is limited to hepatocytes, with periportal hepatocytes (PP-Hep) showing highest expression in normal mice. 9-months post-ALIOS diet feeding, two NASH-specific hepatocyte clusters (mNASH-Hep1 and 2) showed lower *Khk* expression. **(G)** Pseudo time analysis of hepatocyte-specific gene expression through 3 or 9-months post-ALIOS feeding showing that *Khk* expression is downregulated through NASH progression.

Histological assessment confirmed the presence of NASH in the livers of mice subjected to the GAN diet, characterized by substantial accumulation of lipid droplets within hepatocytes, indicating macrovesicular and microvesicular steatosis (Fig. 1-A, top). Immunohistochemical analysis revealed altered expression patterns of ketohexokinase (KHK) in NASH liver tissues compared to healthy liver tissues, with disrupted periportal and interportal distribution (Fig. 1-A, bottom). Western blot analysis of protein lysates from both control and NASH liver tissues confirmed the reduced expression of KHK protein (Fig. 1-C). This was concomitant with the downregulation of fructose transporters, Glut2 and Glut5 but not Hk1 expression (Fig. 1-D).

#### • Spatial and temporal expression of Khk using Single-nucleus RNA sequencing of mice fed with ALIOS diet

To broaden the translational relevance of our findings, we analyzed an additional diet model. The American Lifestyle-Induced Obesity Syndrome (ALIOS) diet^50^, closely mimics Western dietary patterns and provides a highly relevant model for human NASH development. We conducted a secondary analysis of single-nucleus RNA sequencing (snRNA-seq) data from the livers of mice fed the ALIOS diet. This technique offers a detailed examination of cell-type-specific responses to diet-induced obesity. Integrating both GAN and ALIOS diet models will provide a comprehensive understanding of changes in Khk at both the whole-tissue and single-cell levels.

4-week-old male mice were placed on a regular chow or ALIOS diet for 3 or 9 months. These four conditions yielded a total of 28,308 nuclei for snRNA-seq. These cells consisted of both hepatocytes and non-parenchymal cells (NPCs) such as immune, endothelial and stellate cells (Fig. 1-E). KHK transcripts were predominantly expressed by hepatocytes, being more abundant in periportal hepatocytes compared to pericentral or hepatocytes in the intermediate zone. Pseudotime analysis using Monocle2 further confirmed a significant reduction in Khk expression after 9 months on the ALIOS diet, supporting these findings (Fig. 1-F). Altogether, these findings demonstrate that the ALIOS and GAN diets effectively induce NASH in mice, characterized by substantial liver pathology and altered expression of genes involved in fructolysis.

### 2- Hepatic KHK downregulation was also observed in lipolysis-deficient mice fed with normal diet

To determine if KHK can be downregulated in a non-diet-dependent animal model, we selected a genetically engineered mouse model of NAFLD lacking Comparative Gene Identification 58 (CGI58). CGI58 is a hydrolase that acts as a coactivator in lipolysis^51^. Mice lacking CGI58 in the liver (CGI58-KO) display excessive neutral lipid storage and formation of cytosolic lipid droplets^52^. These mice develop steatohepatitis and ultimately, fibrosis under a standard diet. Uniquely, these mice are not insulin resistant, gain weight at the same rate as littermate controls and show no elevation of plasma triglycerides^52^. Therefore, the CGI58-KO model allows investigation of KHK expression in a background of liver inflammation, without the confounding factors of altered insulin resistance or obesity.

We first confirmed the metabolic phenotype of CGI58-KO mice and age-matched controls, at ∼6 months of age, fed standard diet (Fig 2-A). Transcript quantification confirmed lack of CGI58 expression in livers of CGI58-KO mice. As expected, CGI58-KO mice did not display a statistically significant difference in body weight. Serum glucose levels in CGI58-KO trended lower than controls, although this was not statistically significant. Therefore, deficiency of CGI58 in the liver of mice did not result in obesity or hyperglycemia, consistent with results reported in the literature (Fig 2-B).

**Fig (2).**
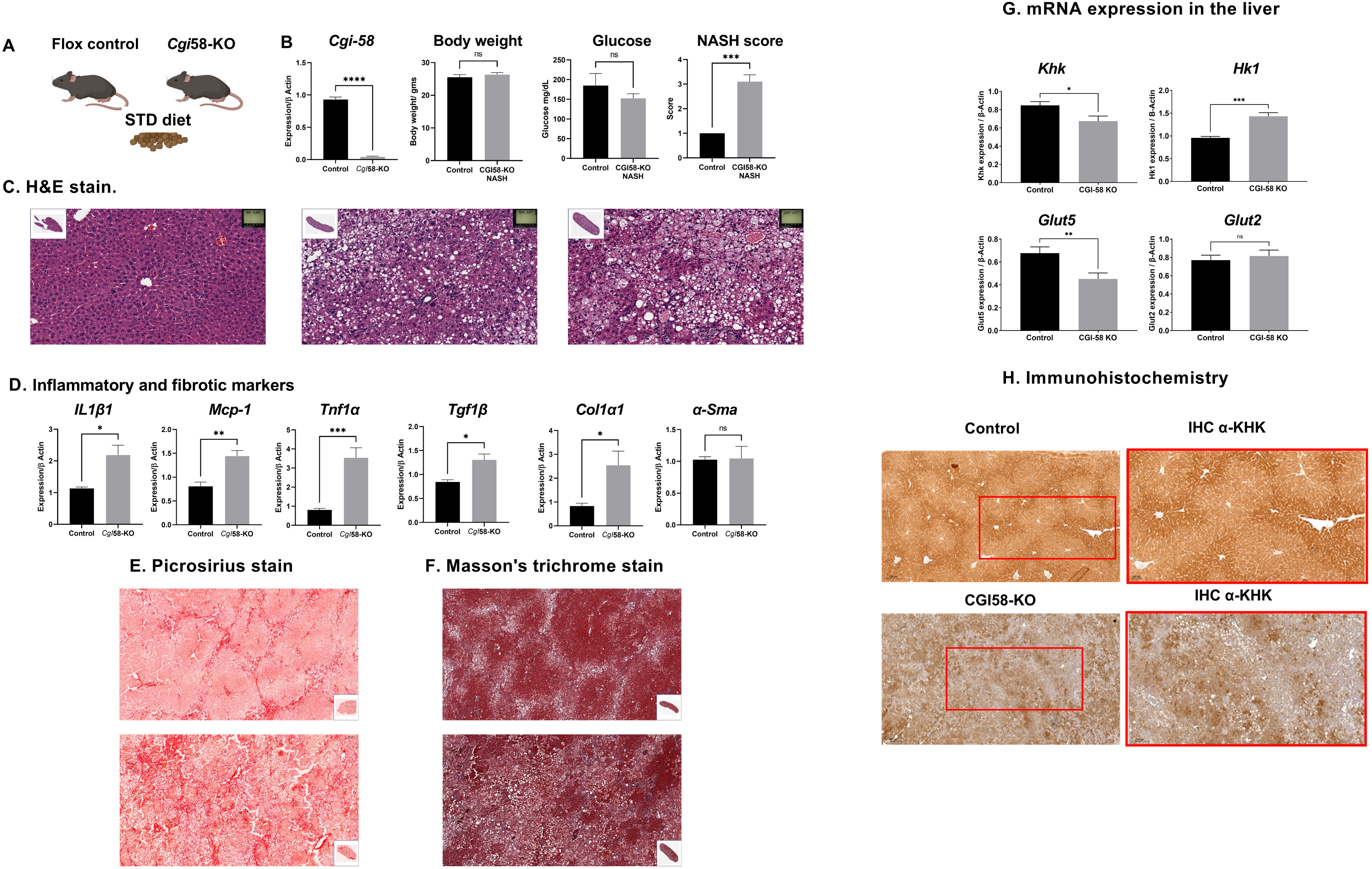
Khk downregulation was associated NASH induced by CGI58 deletion. **(A)** CGI58- KO mice or age-matched controls were fed regular chow. **(B)** CGI58 transcript levels were significantly reduced in CGI58-KO mice but not in controls, but with no change in body weight or blood glucose levels. Histological analysis revealed elevated NASH scores as depicted in **(C)**, representative H&E stains of (left) control or (center and right) CGI58-KO mice. **(D)** Expressions of inflammatory cytokines interleukin 1 beta-1 (*IL1β1*), monocyte chemoattractant protein-1 (*Mcp-1*), tumor necrosis factor alpha (*Tnf1α*), tumor growth factor (*Tgf1β*), were elevated as well as the fibrosis marker collagen 1 alpha 1 (*Col1α1*). Alpha smooth muscle *Actin* trended higher but was not statistically significant. **(E)** Picosirius red and **(F)** Mason’s trichrome staining of two representative sections of CGI58-KO mice revealed fibrosis in these mice. **(G)** Transcript levels of Khk were reduced in CGI58-KO mice, while glucose-specific HK1 increased. Fructose transporter Glut5 reduction mirrored *Khk* reduction, but *Glut2* levels were not significantly altered. **(H)** KHK immunosignal in control livers demonstrated an organized pattern of expression, that showed predominantly periportal expression (red box, inset). Zonal organization was lost in CGI58-KO mice.

However, histological examination of livers in CGI58-KO mice showed obvious signs of disease. Steatosis was clearly observed in these mice, consisting mostly of small vacuoles (microvesicular steatosis) with occasional presence of large fat vacuoles, indicative of macrovesicular steatosis (Fig 2-C). Besides fat accumulation, substantial lobular and periportal inflammation was observed, characterized by robust immune infiltration. Accordingly, pro-inflammatory markers such as Tnfα, Tgf1-ß, Mcp1 and IL1ß were elevated, suggestive of a NASH phenotype (Fig. 2-D).

Markers of fibrosis such as collagen type1α1 (Col-1α1) were significantly elevated but α-smooth muscle actin (α-SMA) did not change significantly although trending higher in CGI58-KO mice, suggesting fibrosis-related hepatocyte remodeling. This is supported by staining with Picrosirius and Masson’s trichrome, where lobular and pericellular fibrosis was observed (Fig. 1-E and F). Collectively, the CGI58-KO model develops liver inflammation and fibrosis in mice fed regular diets, without obesity or elevated blood glucose.

#### KHK expression is reduced in livers CGI58-KO mice

We proceeded to ask if there were differences in KHK expression levels between CGI58-KO, NASH mice and controls. At the transcript level, KHK expression was reduced in CGI58-KO mice compared to controls. Further metabolic reprogramming was observed, where an increase in hexokinase (HK)-1, which favors glucose as a substrate, was elevated in the same mice. At the plasma membrane, glucose transporter (Glut)-5 reduction was observed. Glut-5 has been reported to be selective for fructose, although Glut-2 levels, which is another putative hepatic fructose transporter, were not significantly downregulated (Fig. 2-G).

At the protein level, the normal liver showed strong KHK immunostaining in periportal hepatocytes, extending to interportal zones (Fig. 2-H, top) concomitant with the sn-RNA seq findings. This spatial organization is reminiscent of zonation seen in other metabolic enzymes of the liver. Periportal KHK expression supports the high rates of fructolysis, allowing rapid handling of nutrient-rich, highly oxygenated blood arriving from the intestines through the hepatic portal vein. CGI58-KO mice display similar disorganization of KHK expression (Fig. 2-H, bottom).

Therefore, KHK expression was significantly altered, in terms of expression, structural organization as well as total protein levels.

### 3- Diet-induced NAFLD impacts systemic fructose handling

High-throughput omics technologies, especially metabolomics, have revolutionized biomedical research. It allowed us to obtain an integrated view of the NAFLD/NASH phenotype^53^. This approach has the potential to unveil novel biological insights into disease mechanisms by providing a holistic view of the intricate molecular networks and pathways involved^54^. To understand the metabolomic response to fructose in the plasma, Tibialis anterior (TA) muscle and liver, a targeted stable isotope tracer fate association of 13C fructose was used to analyze metabolic fluxes and flux surrogates with exposure to fructose in control and NASH mice induced by GAN diet.

No significant changes in glycolytic and gluconeogenic pathways between NASH and control groups were observed. However, enrichment analysis showed a significant increase in +3M hexose (at 120 min), lactate (60 and 120 min), and pyruvate levels in the NASH group at various time points (30, 60, and 120 minutes) after injection (Supplementary Fig. 1A). Total alanine levels showed no significant difference between groups. Nevertheless, enrichment analysis of the +3M alanine fraction revealed a significant elevation in the NASH group at 30, 60, and 120 min that mirrors pyruvate (Supplementary Fig. 1B). This suggests an increased contribution of fructose-derived carbons to alanine synthesis, possibly through enhanced gluconeogenesis or transamination reactions. Ketone bodies (3-hydroxybutyrate and acetoacetate) showed a significant elevation in total counts at 90, and 120 mins, but not in the +3M enrichment fraction, in the NASH group (Supplementary Fig. 1C). Such observation indicates an overall increase in ketogenesis in the NASH group, likely due to increased fatty acid oxidation or altered energy metabolism.

### 4- Diet-induced NAFLD reprograms tissue-specific fructose metabolism

The systemic effect of fructose also extended to the target tissues, especially in TA and liver. To delve deeper into these alterations, we investigated the changes in glycolysis / gluconeogenesis, amino acid metabolism and ketogenesis pathways. Total hexose count and lactate was significantly lower in the liver indicating impaired glycolytic activity or increased utilization of pyruvate in the muscle during NASH^55,56^ (Fig.3-A). Yet there was no change in the amino acid metabolism in either TA or liver. Interestingly, total h3-hydroxybuterate levels were significantly lower in the liver (Fig.3-C).

**Fig. (3).**
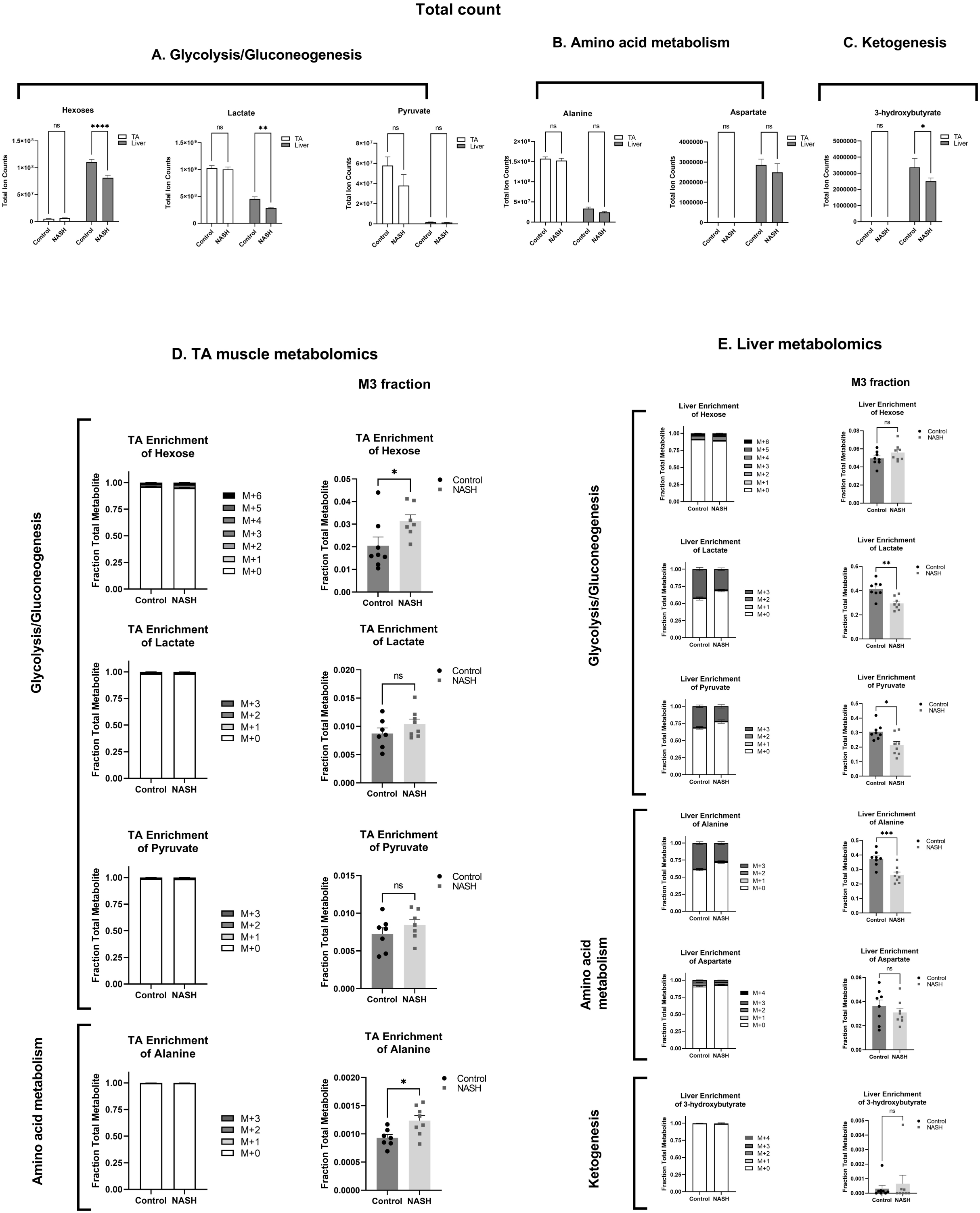
Diet-induced NAFLD reprograms tissue-specific fructose metabolism. (A) Total hexose (glucose and fructose) levels were significantly lower in NASH livers compared to controls. Although no significant differences were observed in hexose enrichment across the tibialis anterior (TA) muscle, liver, and kidney, pyruvate analysis revealed a significant reduction in total levels in the TA muscle. In the NASH liver, the M3 pyruvate fraction was significantly reduced, with a corresponding increase in the M0 fraction, indicating impaired glycolysis and a potential shift towards gluconeogenesis. Lactate levels in the liver were also significantly reduced, with a similar decrease in the M3 fraction and an increase in the M0 fraction, further supporting impaired glycolysis. (B) Alanine levels showed no change in total abundance, but the M3 fraction was significantly reduced in the NASH liver, accompanied by an increase in the M0 fraction. This suggests altered glycolysis and enhanced alanine synthesis through amino acid catabolism or transamination. (C) Ketogenesis, represented by 3-hydroxybutyrate, showed no significant difference in the liver. (D) In the TA muscle, hexose levels were significantly increased in NASH, with no changes in lactate or pyruvate levels. However, the M3 alanine fraction was significantly elevated. (E) In the liver, M3 lactate and pyruvate fractions were significantly reduced, with no changes in hexose levels. Alanine levels were also reduced, while ketone body levels (3-hydroxybutyrate) remained unchanged.

#### • In the TA muscle

The differential enrichment analysis of the TA skeletal muscle, NASH induced the increase in +3M fraction of hexose suggesting increased fructose uptake and breakdown. Concomitantly, no changes were observed in lactate and pyruvate levels between the two groups (Fig.3-D). Moreover, there was a significant increase in fructose-derived alanine synthesis suggesting a metabolic adaptation enabling the muscle to utilize fructose as an energy source, especially under conditions of limited glucose availability^57^. Alanine can be transported to the liver, where it can be converted back to pyruvate and enter gluconeogenesis or the tricarboxylic acid (TCA) cycle, thus contributing to overall energy homeostasis. This process is consistent with the observed increase in plasma +3M enriched alanine levels.

#### • In the liver

The hexose total count in the liver exhibited a significant reduction in NASH compared to the control (Fig. 3-E), potentially indicating a decreased glycolytic flux or impaired glucose metabolism in the liver during NASH progression^35,58^. Concurrently, total lactate levels in the liver also showed a significant reduction in NASH relative to the control, suggesting a reduced conversion of pyruvate to lactate (Fig. 3-E), which may be due to altered lactate dehydrogenase activity or changes in lactate clearance mechanisms^55,56,59^. In the liver of NASH mice, ketogenesis was markedly reduced, as evidenced by decrease of 3-hydroxybutyrate level (Fig. 3-E), suggesting a shift towards an alternative energy source or a compensatory mechanism in response to the metabolic disturbances associated with NASH. Therefore, mice subjected to GAN diet developed NASH, that impaired glycolysis and reduced β-oxidation, particularly in the liver. Although the differential enrichment analysis of +3M hexose levels were unchanged, lactate and pyruvate levels were significantly reduced in the NASH group compared to controls. This effect extended to a reduction in liver alanine levels in the NASH group. Ketone body (3- hydroxybutyrate) levels showed no significant change between the NASH and control groups (Fig. 3-E). The combined reduction in lactate, pyruvate, and alanine levels, along with stable ketone body levels, points towards mitochondrial dysfunction, which is a hallmark of NASH and can lead to impaired energy metabolism, reduced TCA cycle activity, and altered substrate utilization^53,60,61^.

Collectively, mice under GAN diet developed NASH and exhibited altered hexose metabolism, with increased hexose levels in the TA muscle and reduced lactate and pyruvate levels in the liver. Additionally, alanine synthesis from fructose-derived carbons was enhanced in the TA muscle but reduced in the liver of NASH mice. However, ketogenesis, as measured by 3- hydroxybutyrate levels, remained unchanged in the NASH group compared to controls.

### 5- GH-IGF1 axis modulates KHK expression in the liver

These disturbances in fructose metabolism developed by NASH influenced us to delve deeper into the mechanism responsible for modulating KHK, a key enzyme in fructose metabolism. KHK phosphorylates fructose into fructose-1-phosphate in a process lacks negative feedback^20^. Interestingly, knocking out the growth hormone receptor (GHR) was associated with elevated Khk expression^45^.

Deletion of the growth hormone receptor in the liver (GHRLD) results in a fourfold increase in circulating GH levels while reducing IGF1 levels by approximately 90%^44^. These mice displayed insulin resistance, glucose intolerance, elevated levels of circulating free fatty acids, and marked hepatic steatosis^41,42^. Infusing IGF1 into these mice partially restored KHK levels, whereas overexpression of STAT5 returned KHK levels to normal. Hence, the GH-IGF1 axis plays a crucial role in modulating KHK expression in the liver and directly impacts fructose metabolism.

To further investigate this axis, we examined the expression of ketohexokinase (*Khk*) in three transgenic mouse models: mice overexpressing bovine growth hormone (bGH), liver-specific growth hormone receptor knockout (GHRKO) mice, and mice overexpressing the growth hormone antagonist (GHA). Remarkably, *Khk* expression was upregulated in the livers of GHRKO and GHA mice (both models with decreased GH action) but was greatly downregulated in mice overexpressing bovine GH (Fig. 4-A). The downregulation of *Khk* was concomitant with the upregulation of *Igf1* and its receptor *Igf1r*.

**Fig. (4):**
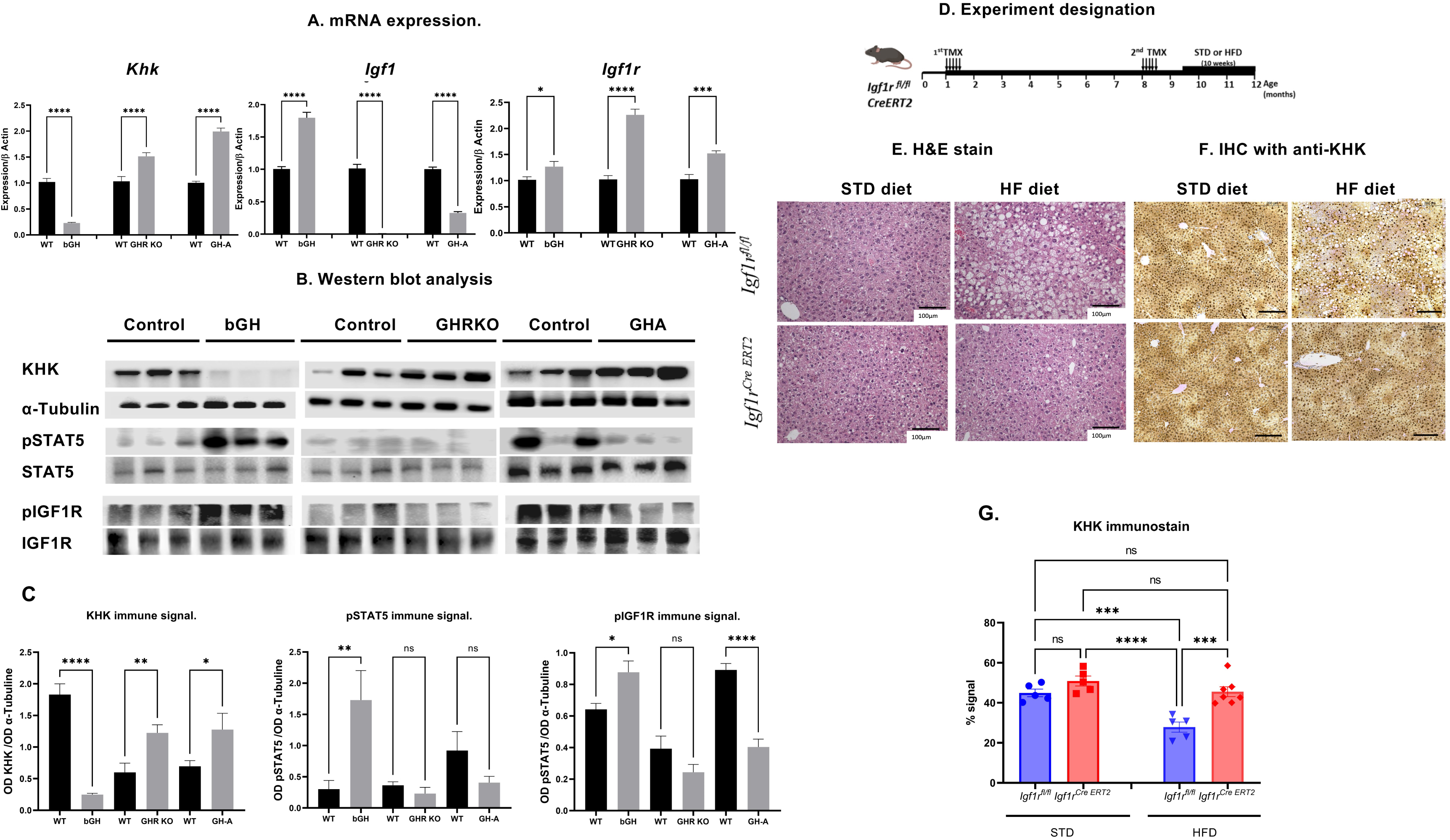
GH-IGF1 axis modulates KHK expression in the liver. (A)qPCR analysis of the expression levels of KHK showed that in bGH, was significantly downregulated while GHRKO and GHA showed a significant upregulation of KHK. This was concomitant with the upregulation of Igf1 expression in bGH but not in GHR and GHA. Interestingly, Igf1r expression was significantly upregulated in all transgenic mice. (B) In bGH, western blot analysis of GH signaling pathway of showing a significant upregulation in pSTAT5. This was associated with the upregulation of pIGF1R and remarkable reduction in KHK immunosignal. On the other hand, knocking out GHR or GHA was concomitant with the absence of pSTAT5 and pIGF1R and a significant upregulation of KHK. (C) Optical density KHK, pSTAT5, and pIGF1R mirrors the qPCR results and shows that KHK expression was directly modulated with GH-IGF1 axis. (D)Experimental designation of the effect of IGF1R deficiency on CreERT2 middle aged mice. Tamoxifen (TMX) was administered daily for 5 consecutive days to *Igf1r^fl/fl^* and UBC- CreERT2; *Igf1r^fl/fl^* four weeks old animals and at eight-months old animals, to generate control *Igf1r^fl/fl^* and IGF1R-deficient CreERT2 mice. Mice were sub divided into two different diet groups: one kept on normal standard diet (STD), and the other changed to a high-fat diet(HFD) and sacrificed on month 12. (E) Representative H&E micrographs of *Igf1r^fl/fl^* under HFD showing macrovesicular and microvesicular steatosis, a hallmark of NASH, top. Conversely, *Igf1r^creERT2^* mice showed only sporadic macrosteatosis, bottom. (F) Immunohistochemical analysis reveals that in *Igf1r^fl/fl^* and *Igf1r^creERT2^* liver tissue under STD, KHK exhibits a periportal and interportal expression pattern. However, when NASH is induced by the high-fat, high-fructose diet in *Igf1r^fl/fl^*, this expression pattern of KHK is disrupted, leading to a significant reduction in KHK immunosignal but not in the liver of *Igf1r^creERT2^*. (G) Quantification of KHK immunosignal showed a significant reduction of KHK when control mice challenged with HFD. This effect was not seen when Igf1r was deleted.

Western blot analysis of liver lysates from these three transgenic mouse models further confirmed these results. There was a significant increase of ketohexokinase (KHK) immunosignal in the GHRKO and GHA groups, while the KHK immunosignal was greatly reduced in the bGH group (Fig. 4-B). These observations align with those seen in liver-specific growth hormone receptor knockout (GHRLD) mice previously published^45^. Elevated KHK levels in the GHA mouse livers were associated with the inactivation of the IGF-1 receptor (IGF1R) signaling pathway (Fig. 4-B), reinforcing the role of the GH-IGF1 axis in regulating KHK and fructose metabolism. Therefore, disrupting growth hormone action, either by GHA overexpression or liver-specific GHR deletion, causes a marked increase in hepatic KHK expression, which correlated with impaired IGF1R signaling in the liver.

### 6- *Igf1r* deletion protects against diet-induced NASH and is associated with increased KHK expression

Thus far, we have demonstrated that GH-IGF1 signaling may play a role in regulating KHK expression in mouse liver. To further investigate the role of IGF1/IGF1R signaling in the regulation of ketohexokinase (KHK) expression, we utilized liver-specific IGF1R knockout mice (*Igf1r^Cre^ ^ERT2^*). Middle-aged (12 months) IGF1R-deficient mice did not exhibit any significant metabolic alterations and appeared normal compared to the control *Igf1r^fl/fl^* mice^62^.

Two groups of 12-month-old *Igf1r^Cre^ ^ERT2^* and control *Igf1r^fl/fl^* were subjected to either standard (STD) or high fat diet (HFD) for 8 weeks to induce non-alcoholic steatohepatitis (NASH) in the liver (Fig.4-D). Histopathological analysis using hematoxylin and eosin (H&E) staining revealed that control *Igf1r^fl/fl^* mice on the high-fat diet developed severe NASH characterized by massive macrosteatosis (large lipid droplet deposition) around the portal spaces. Interestingly, hepatocytes in the pericentral region exhibited microsteatosis with smaller lipid droplet accumulation (Fig.4-E, top). In contrast, the liver-specific *Igf1r^Cre^ ^ERT2^* mice challenged with the high-fat diet displayed only sporadic macrosteatosis (Fig.4-E, bottom). This observation suggests that deletion of the IGF1 receptor protected the liver from the development of diet-induced NASH.

In *Igf1r^fl/fl^* (control) and *Igf1r^CreERT2^* (knockout) mice subjected to standard diet (STD), immunohistochemical analysis revealed strong KHK immunosignals in periportal hepatocytes, extending into the interportal zones, (Fig.4-F, right). However, in *Igf1r^fl/fl^* mice that developed (NASH), immunohistochemical analysis showed a disorganized and reduced KHK immunosignal pattern (Fig.4-F, top left). Surprisingly, *Igf1r^CreERT2^* mice challenged with a high-fat diet did not develop NASH and maintained an intact KHK immunosignal distribution, like the control mice (Fig.4-G). Therefore, while NASH development in *Igf1r^fl/fl^* mice disrupted the normal periportal zonation of KHK expression, *Igf1r^CreERT2^* mice were protected from this KHK disorganization despite high-fat diet challenge.

### 7- GH-IGF1 activation promotes KHK degradation in HepG2 cells

Animal models and primary animal hepatocytes are commonly used to study hepatic lipid metabolism; however, their findings may not always translate effectively to humans due to significant species-specific differences in lipid metabolism^63^. To establish a relevant human liver cellular model for investigating the impact of GH-IGF1 on KHK expression, we utilized a hepatoma cell line, HepG2. These cells can proliferate continuously in the presence of either glucose or fructose as a sole carbon source^37^.

To elucidate how the GH-IGF1 axis regulates ketohexokinase (KHK), we employed HepG2 cells and exposed them to recombinant IGF1 (100 ng/mL) or recombinant human growth hormone (HGH, 10 ng/mL) for 15 minutes in the presence of increasing fructose concentrations (Fig.5-A). IGF1 treatment activated the IGF1 receptor (IGF1R), as indicated by increased phosphorylation at Y1135 on the β-subunit and subsequent activation of AKT through phosphorylation at Ser473 (Fig. 5-B). Notably, IGF2 treatment also activated its receptor and led to KHK degradation, highlighting the role of the IGF axis in KHK regulation (data not shown).

**Fig. (5).**
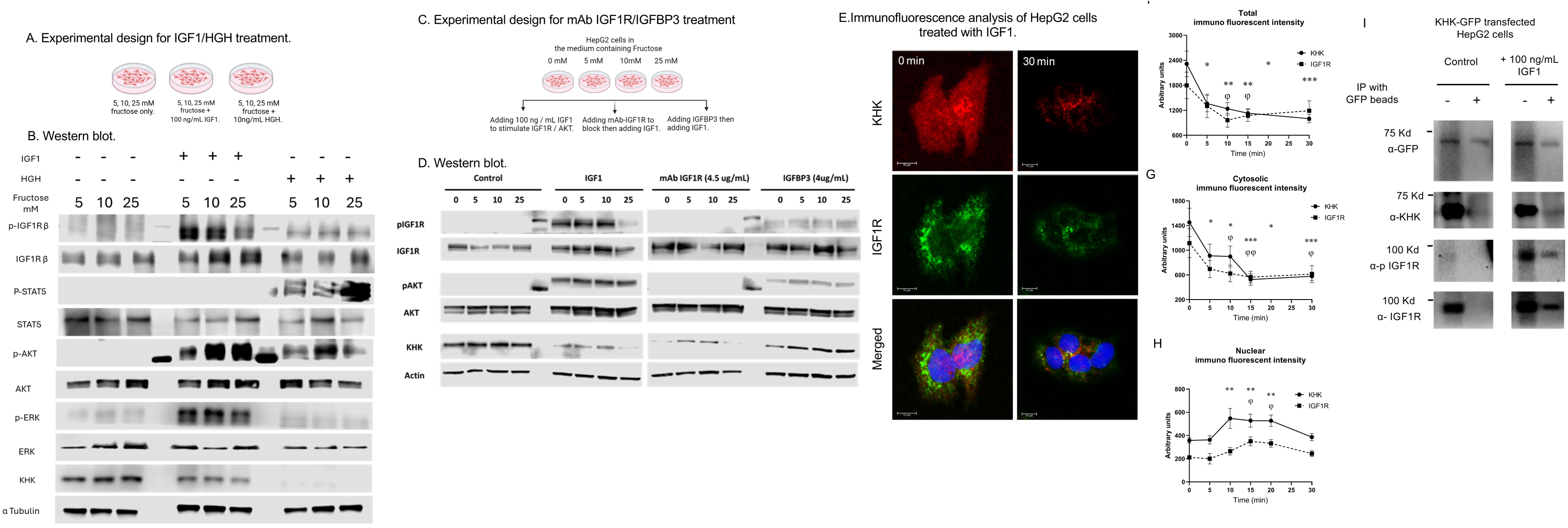
HGH and IGF1 modulate KHK immunosignal in HepG2 cells. (A) graphical summary of the HepG2 treatment. (B) Western blot analysis revealed that treatment with Insulin-like Growth Factor 1 (IGF1) induced the phosphorylation of its receptor, IGF1Rβ, as well as the downstream kinase AKT473. This phosphorylation was associated with a reduction in the immunosignal of Ketohexokinase (KHK). On the other hand, treatment with Human Growth Hormone (HGH) induced the phosphorylation of the downstream transcription factor STAT5. Interestingly, HGH treatment also stimulated the phosphorylation of IGF1Rβ, which was associated with a complete reduction in the KHK immunosignal, (C). Experimental design for inhibition of IGF1R activation. (D) Notably, inhibiting IGF1R with Teprotumumab successfully inhibited receptor phosphorylation but did not rescue the KHK immunosignal. Conversely, treatment with insulin-like growth factor binding protein 3 (IGFBP3) successfully inhibited IGF1R phosphorylation and restored the KHK immunosignal, (B). The STAT5 inhibitor effectively suppressed the phosphorylation of STAT5, consequently restoring the immunosignal of ketohexokinase (KHK), (C). (E) The KHK immunofluorescent signal (red) appeared cytosolic, while the IGF1R signal (green) was observed as cytosolic inclusions near the perinuclear region. (F) The total immunofluorescent signals of both KHK and IGF1R significantly decreased immediately after 5 minutes of IGF1 stimulation. (G) Cross-sectional analysis confirmed that the reduction was predominantly in the cytoplasm. (H) The change in the immunofluorescent signals of both proteins in the nuclear region did not match the reduction in the cytosol; instead, it increased between 10 and 20 minutes. This indicates that the reduction in KHK signal was due to lysosomal degradation rather than nuclear translocation. (I) HepG2 cells were transfected with either GFP (control) or KHK-GFP. Cell lysates were used as inputs (-) or immunoprecipitated (+) with anti-GFP-conjugated beads. KHK-GFP appeared as a single band at 62 kDa when blotted with anti-GFP. However, multiple bands (indicated by arrows) were observed when blotted with anti-KHK. Only phosphorylated IGF1R (p-IGFR) was pulled down when HepG2 cells were treated with IGF1, suggesting that KHK interacts with IGF1R upon its activation.

Interestingly, HGH treatment induced on-target activation of STAT5 at Y694, a downstream marker of GHR signaling. Notably, HGH also resulted in phosphorylation of IGF1R and the complete loss of KHK immunosignal, indicating crosstalk between GH and IGF1 signaling pathways in the regulation of KHK expression.

Given that IGF1R signaling is transduced via the activation of two major kinase pathways, PI3K/AKT and MAPK/ERK ^64^, we explored whether inhibition of either ERK or AKT could significantly protect KHK against degradation. Indeed, HepG2 cells treated with IGF1 effectively induces autophosphorylation of its receptor. Importantly, we found that increasing concentrations of the AKT inhibitor MK-2206 dihydrochloride significantly inhibited AKT phosphorylation and protected against IGF1-induced KHK degradation (Suppl. Fig. 3-A). Additionally, treatment with the ERK1/2 inhibitor U0126 also prevented IGF1-induced KHK degradation (Suppl. Fig.3-B). These findings highlight the crucial roles of both the ERK and AKT signaling cascades in mediating IGF1-induced KHK degradation.

#### • IGFBP3 but not Teprotumumab can restore KHK in HepG2 cells

IGF-mediated signaling is dependent on cognate receptor binding, in competition with associated binding proteins, IGFBPs^64,65^. The half lifetime of IGFs in biological fluids does not exceed a few minutes, but is increased when binding with IGFBPs, through formation of a ternary complex. We asked if addition of IGFBP (Fig.5-C) would result in reduced KHK degradation, through sequestering IGF1 from binding its cognate receptor^66^. Indeed, IGFBP3 resulted in reduced pIGF1R Y1135 phosphorylation, suggesting inhibition of growth factor receptor binding. Downstream signaling via AKT was also reduced, as evidenced by lowered pAKT S473 phosphorylation. Significantly, KHK expression was restored, albeit to levels lower than non-IGF1 treated cells (Fig.5-D).

While IGF1 signaling can be inhibited using growth factor neutralizing antibodies, we found that Teprotumumab, a humanized monoclonal antibody (mAb) recently approved for treatment of Graves’ disease treatment^67^, effectively inhibited phosphorylation of IGF1R and AKT but surprisingly did not restore KHK expression. These findings suggest that while IGFBP3 can sequester IGF1 and prevent its binding to the receptor, thereby reducing downstream signaling and restoring KHK expression, the mechanism by which Teprotumumab inhibits IGF1R signaling doesn’t lead to the same outcome for KHK regulation (Fig.5-D).

Taken together, we have shown that IGF activation results in KHK degradation, through activation of IGF1R. Sequestration of IGF1 using IGFBP3 can restore KHK downregulation, but IGF1R specific mAb does not rescue KHK degradation. These results suggest a distinct mechanistic link between IGF-IGFBP interactions that are not recapitulated using IGF-IGF1R mAb, warranting future investigation.

### 8- KHK degradation is associated with lysosomal vesicles formation upon IGF1/HGH stimulation in HepG2 cells

To this point, we showed that stimulation with IGF1 or HGH was associated with KHK degradation within 15 minutes. Upon GH-IGF1 stimulation, both GHR (Suppl. Fig. 5-A) and IGF1R (Suppl. Fig. 4) undergoes ligand-dependent internalization via both clathrin- and caveolin-mediated endocytic pathways^64,65,68^. Post-internalization, these receptors are either recycled back to the cell surface or degraded through proteasomal and lysosomal pathways^64^, a process that KHK likely plays a crucial role.

To further delve into this mechanism, we incubated HepG2 cells with IGF1 for various time points (0, 5, 10, 15, 20, and 30 minutes). IGF1 treated cells were then fixed and stained with anti-KHK (green) and Lysotracker, a dye used to detect lysosomes (red), (Suppl. Fig. 6-A). In unstimulated cells (0 min), the immunofluorescent signal for KHK was primarily observed in the cytosol.

Interestingly, lysosomal bodies (red) appeared at 5 minutes and continued to increase up to 30 minutes. This increase corresponded with a reduction in the fluorescent signal of KHK. Quantification of the Lysotracker fluorescent signal showed a logarithmic regression over time, (Suppl. Fig.6-B). In contrast, the quantification of KHK immunofluorescence displayed an exponential decay starting immediately after 5 minutes of exposure to IGF1 (Suppl. Fig.6-C).

Double immunofluorescent staining of treated HepG2 cells with anti-KHK and anti-GHR antibodies reconfirmed distinct localization patterns. In unstimulated cells, the KHK signal remained cytosolic, whereas the GHR signal appeared as cytosolic inclusions in the perinuclear region (Suppl. Fig. 5-A). Notably, the total immunofluorescent signals of both KHK and GHR showed a significant reduction immediately following HGH stimulation (Suppl. Fig. 5-B and C). This suggests a rapid cellular response to IGF1, leading to the degradation of these proteins.

Cross-sectional analysis of KHK and IGF1R immunofluorescent signals confirmed that the decrease occurred mainly in the cytoplasm (Fig. 5-E). Interestingly, the nuclear KHK signal did not show the same reduction; instead, it increased between 10 and 20 minutes (Fig. 5 F-H), suggesting that the cytosolic reduction was due to degradation rather than nuclear translocation.

To examine the potential interaction between KHK and IGF1R, HepG2 cells overexpressing KHK-GFP were either left untreated as controls or stimulated with 100 ng of human IGF1. Cell lysates from both conditions were subjected to immunoprecipitation using GFP-catcher, followed by western blot analysis with anti-KHK, anti-phospho-IGF1R, and anti-total IGF1R antibodies. Notably, phosphorylated IGF1R was successfully co-immunoprecipitated with KHK-GFP, indicating that KHK interacts with the activated IGF1R (Fig.5-I).

Collectively, these findings suggest that the rapid degradation of KHK in HepG2 cells upon IGF1 stimulation occurs in the cytosol without translocation to the nucleus.

### 9- Hepatocytes specific upregulation of IGF1R corresponds with KHK downregulation in diet-induced NASH

We investigated whether IGF1R activation is associated with NASH progression by performing a secondary analysis on snRNA-seq data from mice fed with the ALIOS diet. IGF1 expression was ubiquitously expressed across different hepatocyte zones and was also observed in non-parenchymal cells (NPCs) such as macrophages and endothelial cells (Fig. 6-A). Pseudotime analysis using Monocle2 revealed elevated Igf1 levels in hepatocytes, increasing from the basal state through steatosis (3 months on ALIOS diet) and further during NASH (9 months on ALIOS diet) (Fig. 6-B). Igf1r expression remained low during steatosis but dramatically increased in hepatocytes at 9 months post-ALIOS diet, indicating that Igf1r elevation is a late event in NAFLD (Fig. 6-C). Thus, diet-induced NASH elevates Igf1/Igf1r expression, contributing to Khk reduction in the liver.

**Fig. (6).**
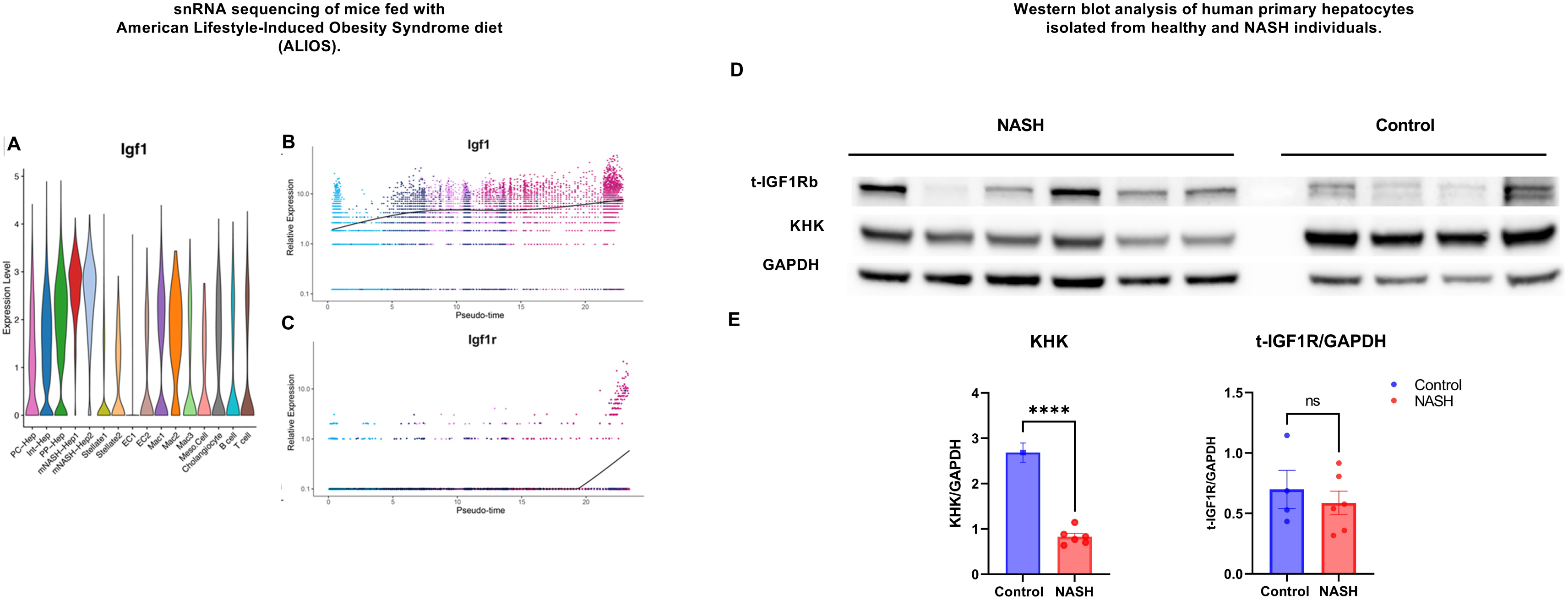
Single nuclear RNA sequencing: **(A)** Violin plots of differential *Igf1* gene expression in mouse livers. *Igf1* expression is limited to hepatocytes, with periportal hepatocytes (PP-Hep) showing low expression in normal mice. 9-months post-ALIOS diet feeding, two NASH-specific hepatocyte clusters (mNASH-Hep1 and 2) showed high *Igf1* expression. **(B&C)** Pseudotime analysis of hepatocyte-specific gene expression through 3 or 9-months post-ALIOS feeding showing that *Igf1/igf1r* expression is upregulated through NASH progression. **(D&E)** Western blot analysis was performed on lysates from cultured primary human hepatocytes isolated from both healthy and NASH individuals. A significant reduction in KHK expression was observed in NASH hepatocytes.

To validate these findings in the context of the GAN diet, we examined liver samples from mice on this diet. Despite the overall reduction in Igf1/2 and their receptors, the ratio of Igf1 to Igf1r was significantly higher in NASH livers compared to controls (Suppl. Fig. 6-A). Primary hepatocytes isolated from control and GAN diet livers showed a significant reduction in KHK immunosignal, associated with an apparent increase in IGF1R phosphorylation (Suppl. Fig. 6-B).

Western blot analysis of primary hepatocytes isolated from healthy humans and patient with NASH further confirmed the downregulation of KHK that was associated with the insignificant reduction in IGF1R expression (Fig. 6-D&E).

Together, these transcriptomic experiments, along with expression analyses, demonstrate the concomitant spatial and temporal expression of KHK and IGF1R in the liver.

## Discussion

Fructose has emerged as a major contributor to the development of NAFLD and its more severe form, non-alcoholic steatohepatitis (NASH) ^2,3^. While glucose is phosphorylated by hexokinase, a rate-limiting enzyme subject to negative feedback regulation, fructose is rapidly phosphorylated by ketohexokinase (KHK), an enzyme that lacks this regulatory mechanism^37^. This unrestricted metabolism of fructose can lead to a rapid depletion of ATP and an accumulation of lipids in the liver, contributing to the development of NAFLD and NASH^21^.

Given the critical role of KHK in fructose metabolism, KHK inhibitors has recently been considered as a potential treatment for NAFLD/NASH^69–75^. Several studies have attempted to identify factors that regulate KHK expression. For instance, negative regulators of KHK has been reported via HIF-2A and peroxisome-deficiency^76^. Conversely, positive regulators of KHK activity, such as carbohydrate response element-binding protein (CHREBP), binds to carbohydrate response elements (ChoREs) in the promoter region of the KHK gene^77,78^. Hence, understanding these regulatory mechanisms is crucial for clarifying KHK’s role in liver function.

Notably, the immunohistochemical localization of KHK showed its presence in the periportal and interportal zones, areas rich in oxygen and active in oxidative energy metabolism, gluconeogenesis, ureagenesis, and bile formation^63^. However, in NASH, liver zonation is disrupted, leading to perturbations in the spatial expression of genes involved in metabolic zonation^79,80^. This disruption was consistent with our findings and others^76^ that KHK expression is significantly reduced during NASH. Contradictorily, a recent preprint^73^ reported an increase in KHK in NASH, prompting us to investigate and resolve these conflicting results.

To address this, we employed two animal models, the GAN diet^48^ and Cgi58 KO^52^. Both clearly demonstrate NASH in the liver through different mechanisms. Consistent with our findings, livers with NASH showed a significant downregulation in KHK expression as previously reported^37^ and is associated with elevation in pro-inflammatory cytokines expression^48,81^ and extracellular matrix deposition^81,82^. Moreover, the expression of Glut2 and Glut5, the main fructose transporters in the liver, were also reduced. Interestingly, sn-RNA sequencing of KHK in the liver of mice subjected to ALIOS (like GAN diet) confirmed the expression of KHK in the periportal and interportal zones and showed a marked reduction with NASH progression. Collectively, NASH shows apparent inflammation associated with disruption of frucotlytic pathway that includes Khk and fructose transporters in the liver.

The impaired ability of the liver to metabolize fructose in NASH mice was evident in their plasma. Temporal enrichment analysis of the +3M fraction following a 13C fructose injection revealed hexose intolerance in NASH mice. Notably, the levels of pyruvate, lactate, and alanine were elevated, indicating metabolic stress and the subsequent accumulation of these metabolites in the blood like other diet induced NASH models^83,84^.

Moreover, reduction of total hexose and lactate counts in the liver indicate impaired glycolytic flux^85^. Additionally, elevated levels of the +3M fraction of hexose and alanine in the liver are indicative of impaired fructolytic activity, leading to the reduction of glycolytic intermediates and altered amino acid metabolism^48,86^. This metabolic dysregulation is characteristic of conditions like NASH, where hepatic metabolism is significantly altered, contributing to the disease’s pathophysiology^35,86,87^.

These observations led us to explore the possible mechanisms that control KHK expression. Intriguingly, mice with a knock out of the growth hormone receptor (GHR) in mice^88^ as well as humans with GHR mutation, Laron syndrome (LS)^47^, develop hepatic steatosis that is associated with elevated Khk expression^45^. Similarly, we observed a significant increase in ketohexokinase (KHK) in mice overexpressing GH antagonist and liver-specific knockout of the GH receptor (GHR) that was associated with NASH progression^88,89^. Conversely, KHK reduction was observed when overexpressing bovine growth hormone (GH), which coincided with the elevation and activation of STAT5b and IGF1/IGF1R. Notably, GHR deletion was associated with a 90% reduction in circulating IGF1 levels^44^. Our sn-RNA sequencing data from ALIOS mice with advanced NASH further supported the inverse correlation between KHK expression and IGF1/IGF1R expression in the liver.

These findings suggest that targeting KHK to inhibit steatosis could be beneficial in early stages of NASH and in NASH induced by GHR mutations, such as in Laron syndrome (LS)^47,89^. However, the question arises: could blocking the IGF1R receptor in hepatocytes halt the progression of inflammation and NASH in the liver? Interestingly, liver-specific knockout of IGF1R protected the liver against NASH and maintained KHK expression in middle-aged mice.

To investigate the effect of GH/IGF1 on KHK expression, we used HepG2 cells, a hepatoblastoma line known for its fructose avidity^37^. The rapid degradation of KHK within 15 minutes of GH/IGF1 stimulation suggests KHK involvement in receptor-mediated endocytosis and degradation. Typically, activated GHR^68^ and IGF1R^66^ undergo endocytosis in clathrin- or caveolin-coated pits, which are subsequently subjected to proteasomal or lysosomal degradation. Interestingly, Teprotumumab, an IGF1R monoclonal antibody, and IGFBP3 are involved in the regulation of IGF1R signaling^67,90^. Teprotumumab directly targets and inhibits IGF1R, while IGFBP3 modulates IGF1R activity indirectly through its binding to IGF1. Conversely, IGFBP3 administration successfully restored KHK levels and decreased IGF1R phosphorylation.

Surprisingly, we noticed the autoactivation of IGF1R in Huh7 cells, which was associated with a reduction in the KHK immune signal. Autoactivation of IGF1R has been observed in several cell lines and in almost 33% of hepatocellular carcinoma (HCC) cases^91^. This further supports the role of IGF1R activation in KHK degradation and the loss of fructolytic ability in NASH and HCC^37^. Therefore, autoactivation; in Huh7 or exogenous stimulation; in HepG2, of IGF1R drives KHK degradation suggesting its involvement in receptor-mediated clearance. This also explains how HCC loses its ability to metabolize fructose.

In summary, these findings highlight the complex regulation of KHK in the liver, influenced by GH/IGF1 axis in NASH. Understanding these regulatory mechanisms could pave the way for targeted therapies to mitigate NASH progression.

## RESOURCE AVAILABILITY

### Lead contact

Further information and requests for resources should be directed to and will be fulfilled by the lead contact, (Salaheldeen Elsaid)

### Materials availability

This study did not generate new unique reagents.

### Data and code availability

- All data reported in this paper will be shared by the lead contact upon request.
- This paper does not report original code.
- Any additional information required to reanalyze the data reported in this paper is available from the lead contact upon request.

## Methods

### - Cell culture

Either HepG2 or Huh7 cells Dulbecco’s modified Eagle’s medium (25mmol/l glucose), supplemented with 10% (vol/vol) FBS, 1% penicillin, and streptomycin, and 1% Gultamax. Medium was exchanged every 3 days, and cells were trypsinized and reseeded at 1:4 dilution when 80-90% confluence was reached.

For fructose stimulation, cells were starved overnight in serum free high glucose medium then stimulated with 0-, 5-, 10-, and 25-mM fructose for 15 min to investigate the change in KHK immunosignal upon fructose stimulation. On the other hand, cells were incubated for 24 hr in previously mentioned fructose concentrations for gene expression analysis. In some experiments, cells were grown to 80-90% confluency in 10% FBS high glucose DMEM then starved for 24 hrs in serum free medium. After 24 hours, the cells were incubated in serum free medium or with 10 % FBS supplemented with 5-, 10-, and 25-mM fructose. RNA was extracted from cells and cDNA were synthesized to investigate the changes in gene expression responsible for fructose uptake and metabolism.

#### - RNA extraction and qPCR

Total RNA was extracted from 2×10^6^ HepG2 or Huh7 cells using PureLink RNA Mini Kit (Invitrogen) according to manufacturer instructions. iScript™ cDNA Synthesis Kit (Biorad) was used to convert one microgram of high purity RNA into cDNA in a total reaction volume of 20 µL. The rection volume was diluted to 200 uL to reach (5 ng/ µL) .2 µL was used for qPCR (10 ng) per reaction. All materials are indicated in Table (1).

**Table 1.** Materials.

| REAGENT OR RESOURCE | SOURCE | IDENTIFIER |
| --- | --- | --- |
| Chemicals, peptides, and recombinant proteins |  |  |
| PureLink RNA Mini Kit | Invitrogen | 12183018A |
| iScript™ cDNA Synthesis Kit | Biorad | 1708840 |
| SYBR™ Select Master Mix | Invitrogen | 4472908 |
| MicroAmp Optical 96-Well | Invitrogen | N8010560 |
| SurePAGE, Bis-Tris, 4-12%,10 wells | Genscript | M00653 |
| Picosirius red | Electron Microscopy Sciences | 26357-02 |
| Recombinant Human Igfbp-3 | R&D systems | UN675-B3 |
| Recombinant Human IGF-I | R&D systems | 291-G1 |
| Recombinant Human IGF-II | R&D systems | 292-G2 |
| Teprotumumab Recombinant mAb | Invitrogen | MA5-41915 |
| PhosSTOP | Roche | 4906845001 |
| Superblock buffer | Invitrogen | 37535 |
| 4-12% Bis-Tris gel | SurePAGE | M00653 |
| GFP-Catcher beads | Antibodies online | ABIN5311508 |
| Vectashield with DAPI | Vector laboratory | H-1200-10 |
| Antibodies |  |  |
| Anti KHK | Sigma | HPA007040 |
| Anti-pIGF1Rβ (Tyr1135/1136) | Cell signaling | 2250 |
| Anti pan IGF1R β (D23H3) (Rb) | Cell signaling | 9750 |
| Anti-pStat5 (Tyr694) | Cell signaling | 8178 |
| Anti-Stat5 | Cell signaling | 3873 |
| Anti phospho AKT (S473) (Rb) | Cell signaling | 9271 |
| Anti pan AKT (11E7) (Rb) | Cell signaling | 4685 |
| Anti-phospho-ERK1/2 | Cell signaling | 9750 |
| Anti-ERK1/2 | Cell signaling | 4060 |
| Anti- α tubulin | Cell signaling | 14179 |
| Anti β-Actin (Rb) | Cell signaling | 2250 |
| Experimental models: Organisms/strain, Cell lines |  |  |
| Male C57BL/6 mice (Flox) | Animal facility of University | N/A |
| Male C57BL/6 mice (CGI-58 KO) | of Maryland Baltimore |  |
| Software |  |  |
| Prism 9.0 | GraphPad | https://www.graphpad.com/ |

Two microliter (2 µL ∼10 ng of cDNA) of this cDNA was used for subsequent PCR amplification with SYBR™ Select Master Mix (cat. 4472908, Invitrogen) in QuantStudio 5 (ThermoFisher scientific) using the following specific primers for human, Table (2) or for mice, Table (3) were purchased from Integrated DNA Technology, IDT.

**Table 2.**
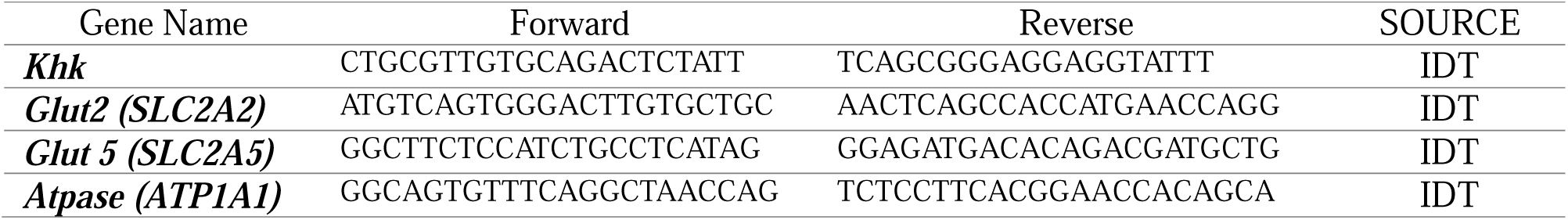
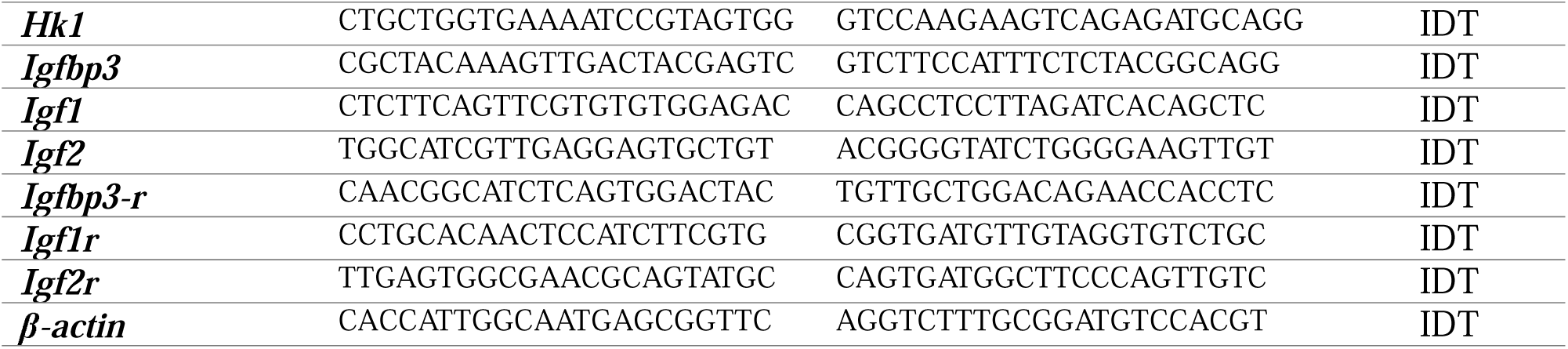
Human (*Homo sapiens*) primers.

**Table 3.**
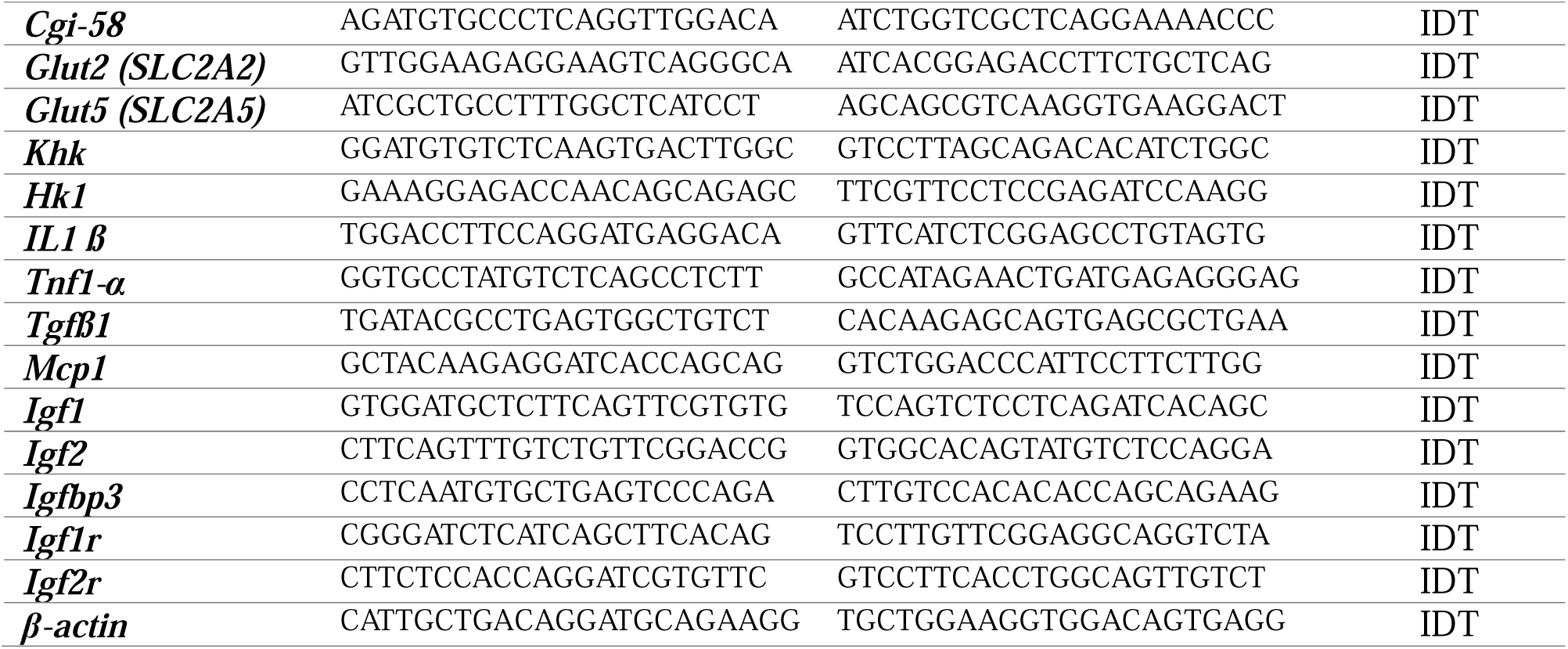
Mouse (*Mus musculus*) primers.

##### - Western blot analysis

Cells were lysed in RIPA buffer, 150 mM NaCl,50 mM HEPES, pH 7.6, containing 0.1% of (100x Protease inhibitor) and one tablet of PhosSTOP (No. 4906845001). Equal amounts of protein samples were blocked in 4X loading buffer and run in 4-12% Bis-Tris gel then transferred to a PVDF membrane. After blocking, membranes were immunoprobed with specific antibodies indicated in Table (1).

### - Immunoprecipitation Using Anti-GFP Agarose Beads

HepG2 cells were transfected with either GFP (as a control) or human KHK-GFP. After transfection, the cells were treated with 100 ng of h- IGF1 for 15 minutes. The cells were lysed and incubated with GFP-catcher beads following the manufacturer’s protocol. The resulting cell lysates and immunoprecipitated proteins were analyzed through western blotting.

### - Immunofluorescence Analysis

Cells were grown on coverslips until they reached 70–80% confluency and then fixed with 4% paraformaldehyde. The cells were permeabilized using PBS containing 0.3% Triton and blocked for 1 hour. They were incubated overnight with the specified primary antibodies, followed by a 1-hour incubation with secondary antibodies. Finally, the cells were mounted with Vectashield and examined under a Leica fluorescence microscope.

### - Mice and diets

Statement of ethics of animal care and use. All animal procedures were approved by the Institutional Animal Care and Use Committees (IACUC) at University of Maryland Baltimore. All animal procedures were carried out in accordance with the U.S. National Institute of Health guidelines, including “Principles for Use of Animals” and “Guide for the Care and Use of Laboratory Animals”. CGI-58 LivKO^7^, bGH^92^, GHRKO^93^, GHA^94^, and Liver-IGF1R KO^62^ mice were generated as described. Homozygous transgene and homozygous floxed mice (Control) were fed ad libitum a standard diet or high fat diet as required after weaning.

### - Liver histopathology

At the end of the experiment, male mice were dissected, and liver specimens were fixed in the 10% buffered formalin solution.

Subsequently, these specimens underwent processing for hematoxylin and eosin (H&E) staining. In addition, paraffin sections were subjected to Picrosirius red staining (Electron Microscopy Sciences, cat# 26357-02) and Mason’s trichrome staining. These staining techniques were employed to detect the presence of extracellular matrix (ECM) deposition and the development of fibrosis.

Additionally, immunohistochemical detection of KHK and IGF1R in liver sections were done with anti KHK antibody (SIG-HPA007040) and anti-IGF1 Receptor β (D23H3) Rabbit mAb (No.9750).

### - Transcriptomic changes and cell clustering of snRNA-seq data

snRNA-seq, snATAC-seq and bulk RNA-seq libraries were sequenced on Illumina NextSeq5000 or NovaSeq. 400 million paired-end reads were obtained for each sample using the sequencing parameters recommended in 10X Genomics guide as described previously^22^.

### - Statistical analysis

Data were analyzed with GraphPad Prism 9 software (GraphPad Software, USA) and presented as Mean ±SEM. We used one-way ANOVA followed by Post HOC Tukey test for multiple comparison. Statistical significance is indicated as figures using the following denotations, \**P*<0.05, \*\**P*<0.01, \*\*\**P*<0.001, \*\*\*\**P*<0.0001.

