## Supplemental Figures for "Hepatic fructose metabolism is antagonized by growth hormone/insulin-like growth factor signaling via regulation of ketohexokinase expression"

### Supplementary Fig. 1- Diet-induced NAFLD impacts systemic fructose handling

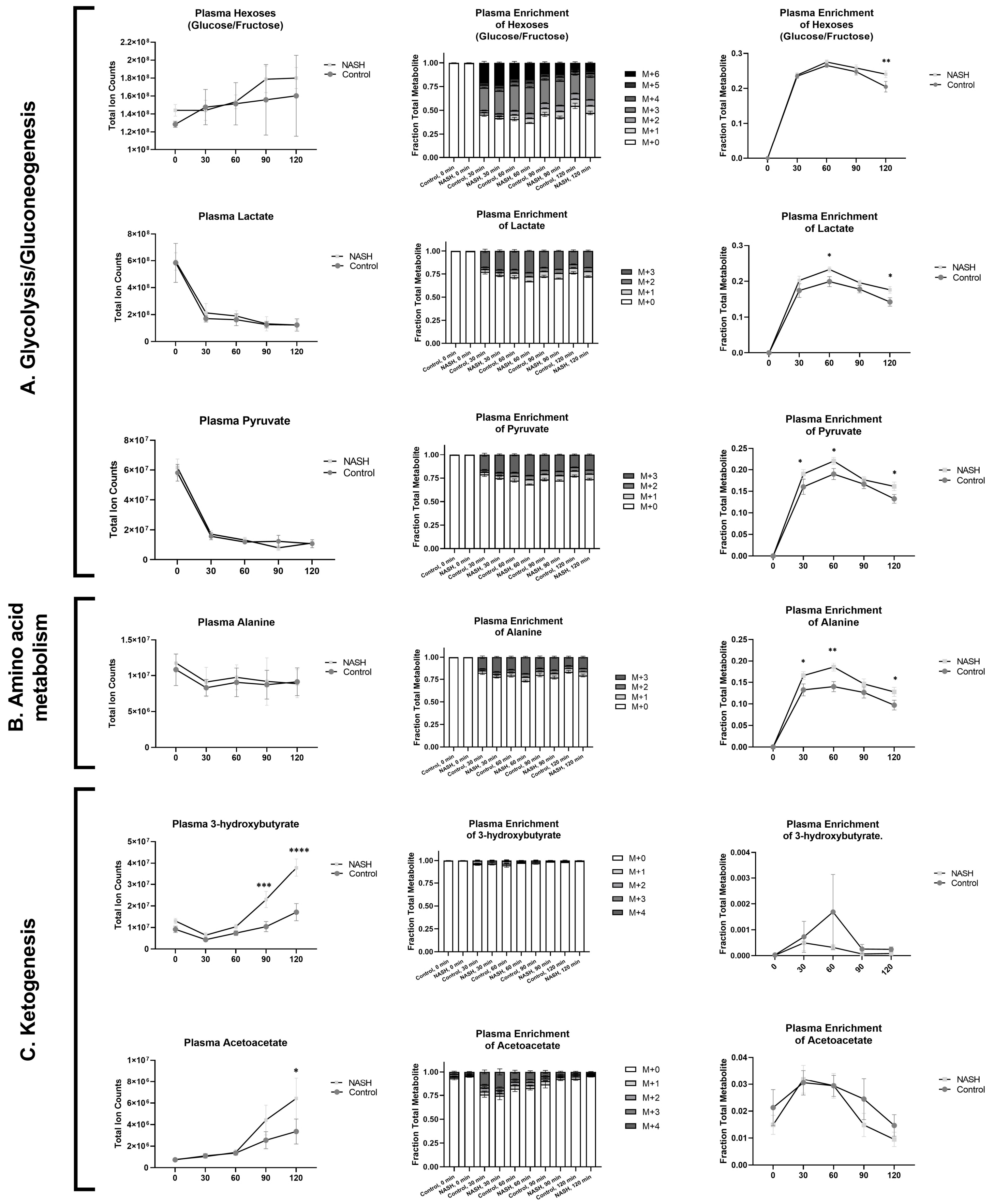

### Supplementary Fig. 2. HepG2 and Huh7, hepatoma vs hepatocellular carcinoma (HCC), display differential response to fructose.

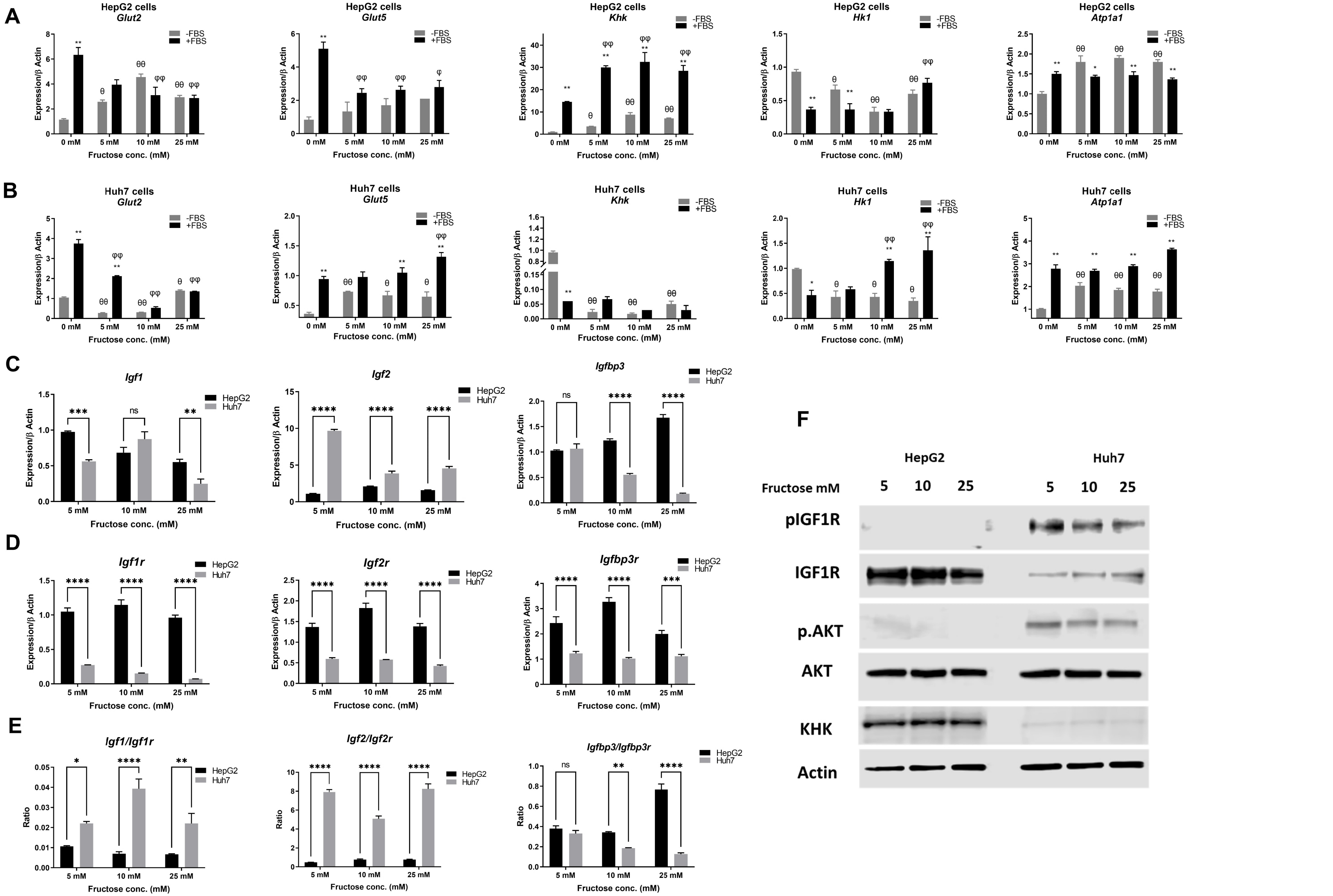

**Supplementary Fig. 3: Effect of ERK AKT inhibitor on restoration of KHK immunosignal in IGF1 stimulation HepG2.**

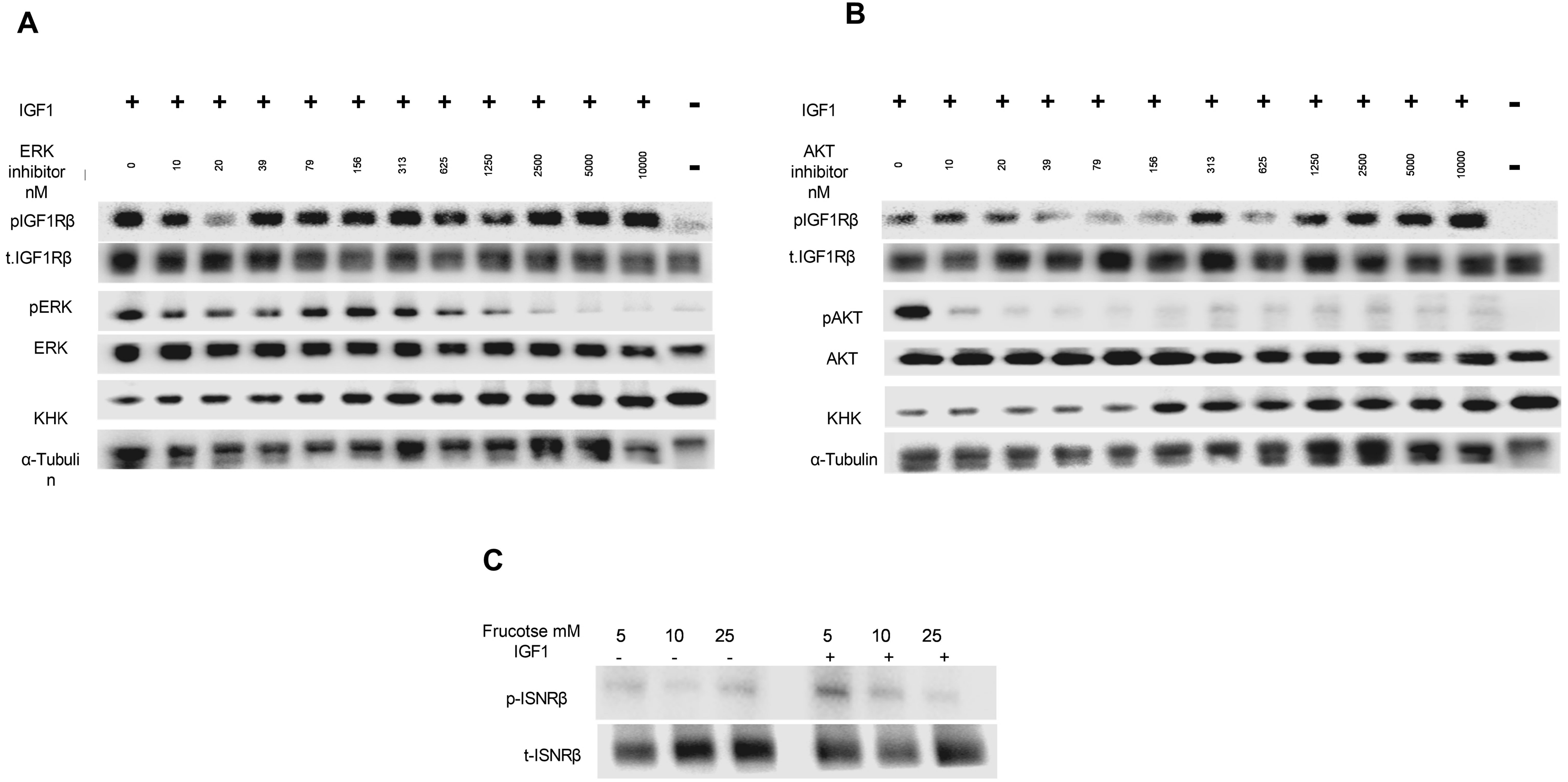

### Supplementary Fig. 4: IGF1 induces IGF1R and KHK degradation in HepG2

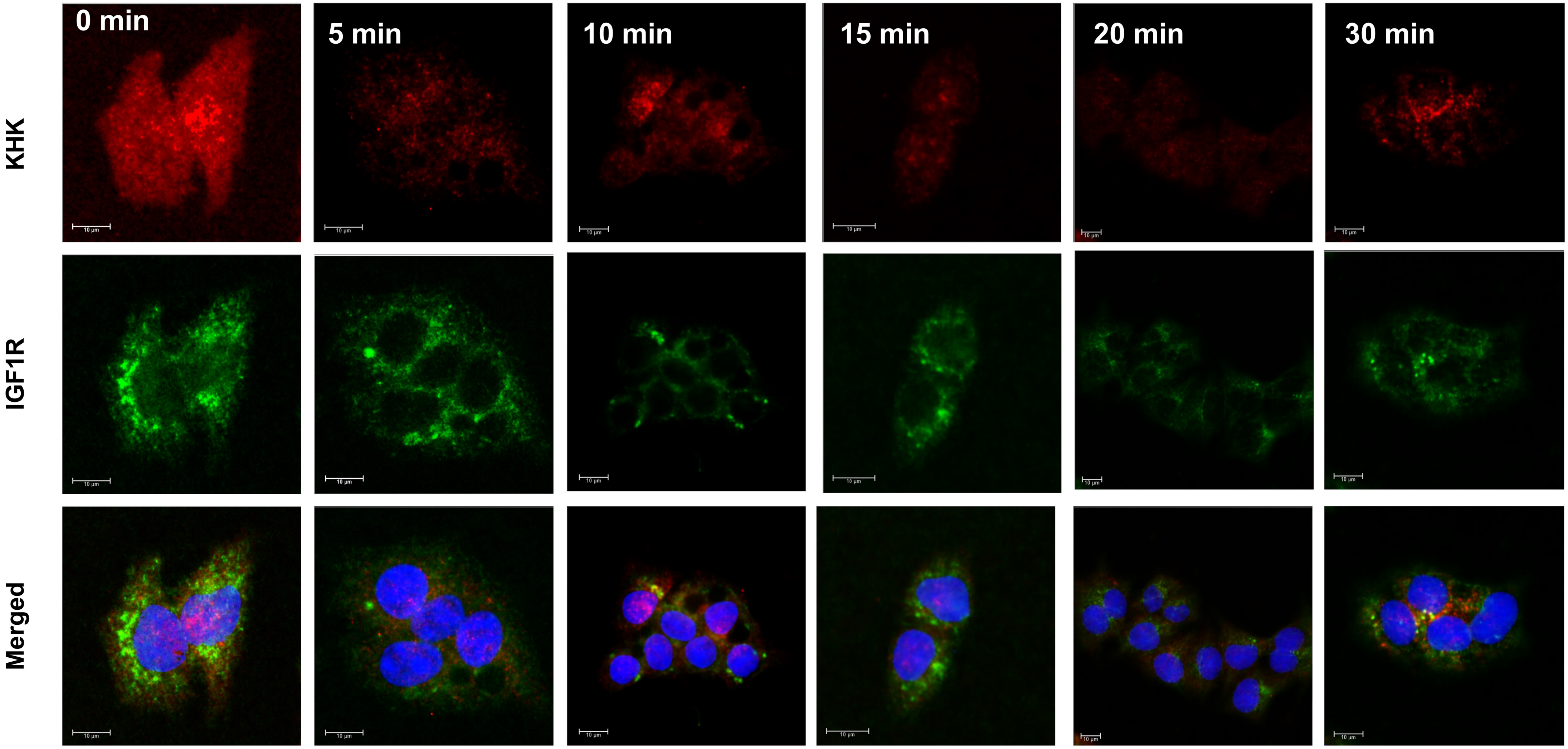

**Supplementary Fig. 5. HGH induces GHR and KHK degradation in HepG2.**

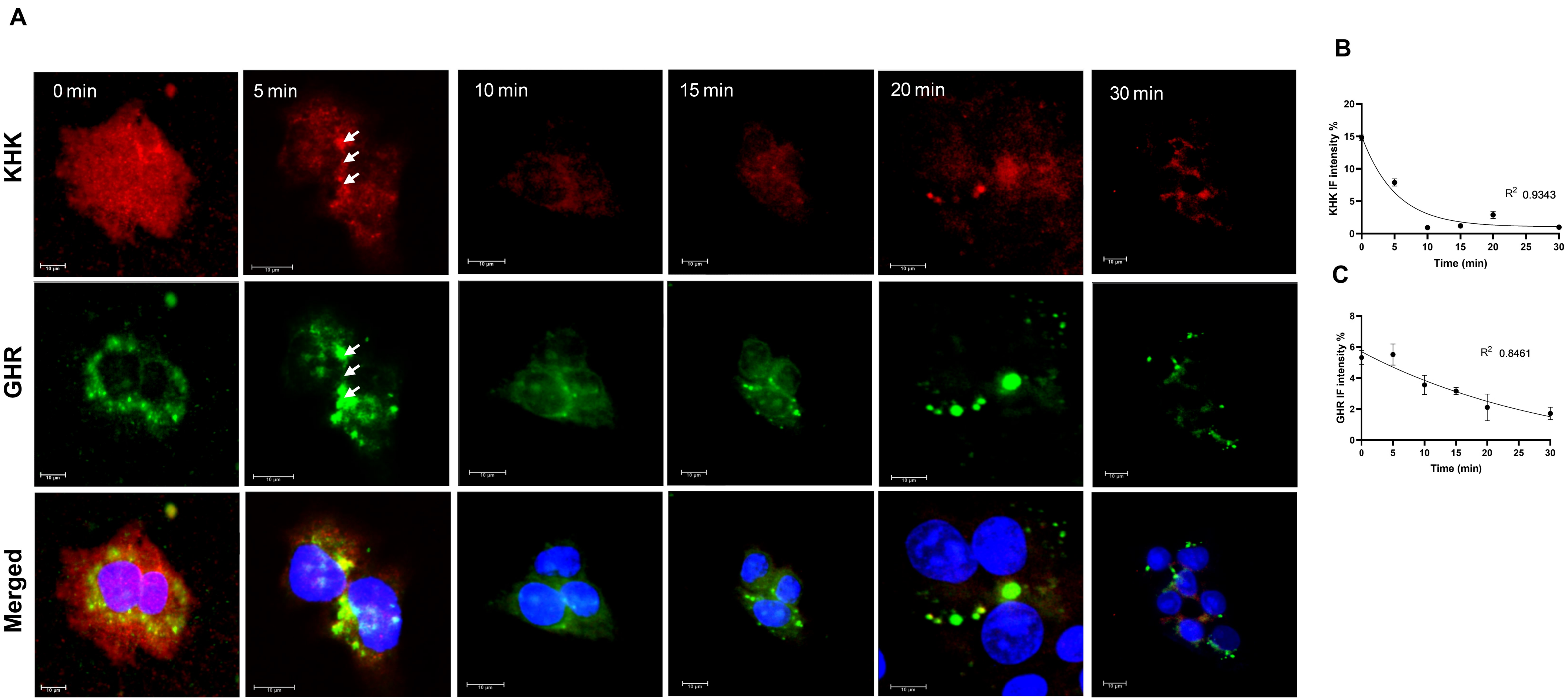

#### Supplementary Fig. 6. IGF1 Induces lysosomal formation and KHK degradation in HepG2

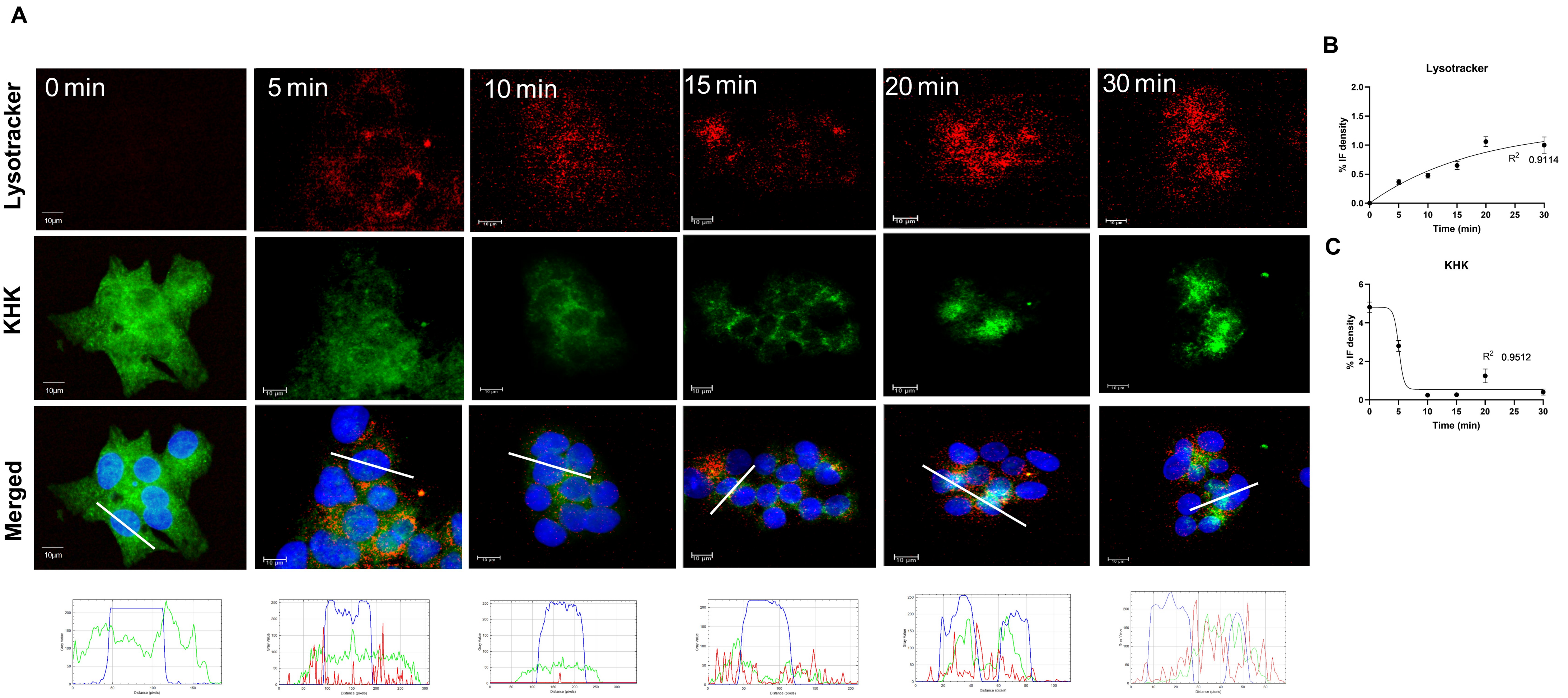

**Supplementary Fig. 7. NASH induced by either *Cgi*-58 knockout (KO) or a GAN diet, activates IGF1 signaling in the liver, leading to KHK reduction.**

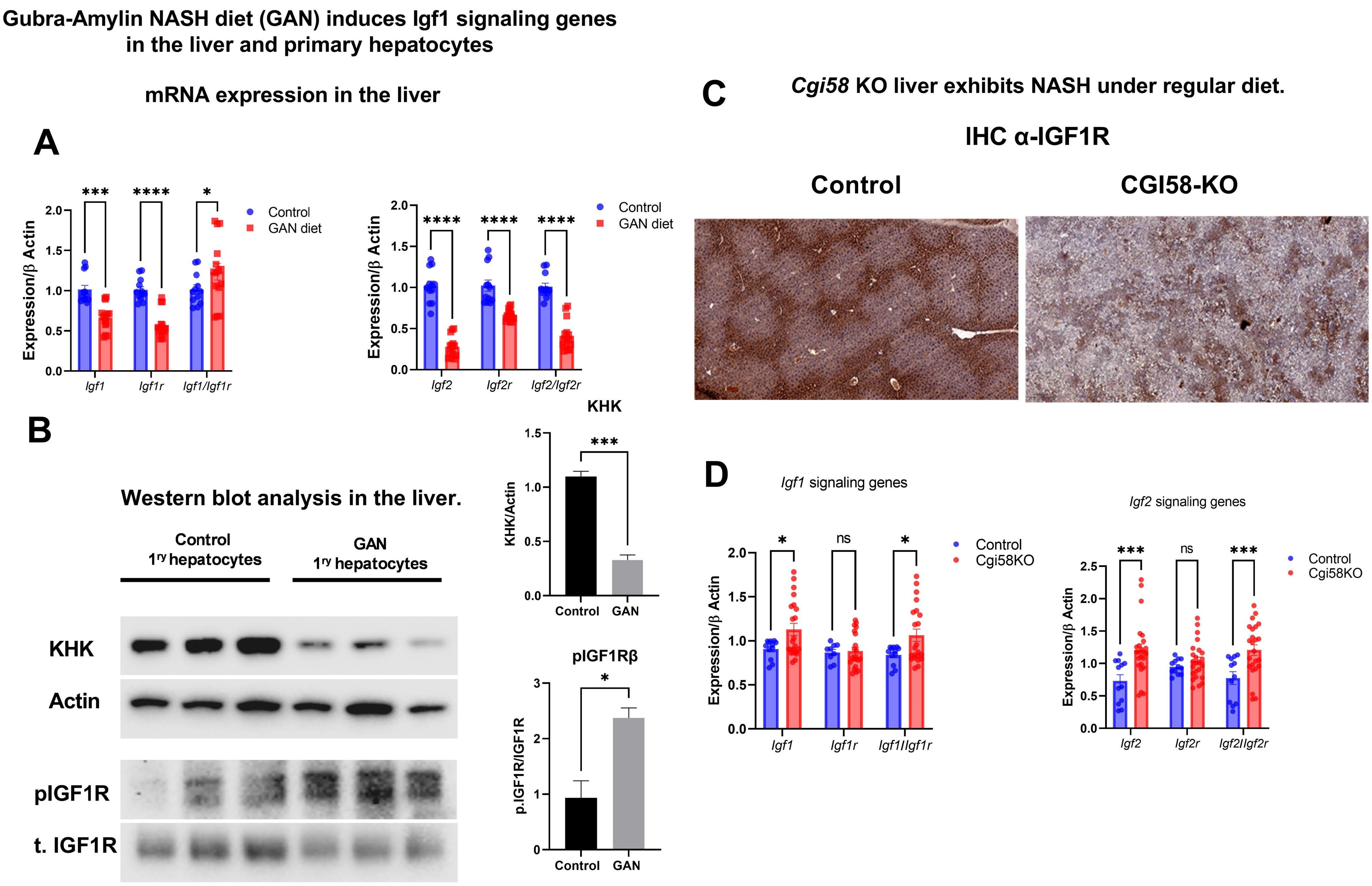
