## Supplemental legends for "Hepatic fructose metabolism is antagonized by growth hormone/insulin-like growth factor signaling via regulation of ketohexokinase expression"

**Suppl. Fig. (1):** **Diet-induced NAFLD impact systemic fructose handling.** (A) Hexose metabolism: No significant changes in glycolytic and gluconeogenic pathways between NASH and control groups. Enrichment analysis showed a significant increase in +3M hexose (at 120 min), lactate (60 and 120 min), and pyruvate levels in the NASH group at various time points (30, 60, and 120 minutes) after injection. (B)Total alanine levels showed no significant difference between groups. However, enrichment analysis of the +3M alanine fraction revealed a significant elevation in the NASH group at 30, 60, and 120 min that mirrors pyruvate. This suggests an increased contribution of fructose-derived carbons to alanine synthesis, possibly through enhanced gluconeogenesis or transamination reactions. (C) Ketogenesis: Ketone bodies (3-hydroxybutyrate and acetoacetate) showed a significant elevation in total counts at 90, and 120 mins, but not in the +3M enrichment fraction, in the NASH group.

**Suppl. Fig. (2):** The expression of the hexose transporters, and enzymes in **(A)** HepG2; and **(B)** Huh7 cells; with fructose as the sole source of carbon +/- 10% FBS. The expression of (C) Khk and Hk1 in HepG2 and (C-E) Comparison of Igf1/2 and Igfbp3 expression between HepG2 and Huh7 cells at varying fructose concentrations.(G): Expression of their receptors in both cell lines under different fructose conditions. (F) Western blot analysis reveals the activation of IGF1R and its downstream kinase AKT in Huh7 cells, accompanied by a significant reduction in KHK levels.

*, significant when comparing -/+ FBS. θ significant when comparing to 0mM fructose -FBS Φ significant when compared to 0 mM fructose +FBS.

**Suppl. Fig. (3):** ERK/AKT inhibitors protects KHK degradation in HepG2 cells. **(A)** IGF1 stimulation successfully activated its receptor and ERK phosphorylation, which was accompanied by KHK degradation. However, incubation with the ERK inhibitor restored KHK immunosignal in a dose-dependent manner.**(B)** While the AKT inhibitor reduced AKT phosphorylation at 10nM, the restoration of KHK did not appear until 313nM. Interestingly, this highlights the different efficacies and thresholds of these inhibitors in protecting KHK from degradation.

**Suppl. Fig. (4):** HepG2 cells were stimulated with IGF1 for 30 minutes and analyzed using double immunofluorescence staining with anti-IGF1R (green) and anti-KHK (red). KHK was distributed throughout the cell without specific subcellular localization, while IGF1R formed cytosolic inclusions. Both signals were noticeably reduced at 5, 10, 15, 20, and 30 minutes post-IGF1 stimulation, indicating a dynamic response to the IGF1 treatment.

**Suppl. Fig. (5):** Double immunofluorescent staining of KHK and GHR in HGH-treated HepG2 cells. (A) The KHK immunofluorescent signal (red) appeared diffused in the cytosol, while the GHR signal (green) was observed as cytosolic inclusions near the perinuclear region. (B) The total immunofluorescent signals of both KHK significantly decreased immediately after 5 minutes of HGH stimulation. (C) Total immunofluorescent signals of both GHR significant decay after 5 minutes of HGH stimulation and extends to 30 min.

**Suppl. Fig. (6):** IGF1 induces spatial and temporal modulation of KHK in HepG2.  (A)The immunofluorescent signal of KHK appears mainly in the cytosol (green). Stimulation with IGF1 (100ng/mL)  induced the formation of lysosomal bodies associated with the reduction in fluorescent signal of KHK. (B) lysotracker (red) was used to detect the lysosomal formation and its quantification showed a logarithmic regression at different time intervals. (C) On the other hand, the regression of quantified KHK immunofluorescence showed an exponential decay immediately after 5 mins of exposure to IGF1.

**Suppl. Fig. (7).** (A) Despite a decrease in Igf1 mRNA in GAN diet-fed mice, ratiometric analysis indicated increased IGF1 activation. (B) Western blot of primary hepatocytes from GAN-fed mice confirmed IGF1Rβ activation, accompanied by a significant reduction in KHK levels. (C) Immunohistochemistry (IHC) of control livers showed a spatial pattern of IGF1R similar to KHK, which was lost in Cgi58 KO livers. (D) Although Igf-1 and -2 expression was elevated in Cgi58 KO livers, receptor levels remained unchanged, with significant ratiometric differences compared to controls.
